# Neural subpopulations in marmoset area MT/MTC detect and discount saccade-related retinal motion

**DOI:** 10.64898/2026.09.04.749428

**Authors:** Amy Bucklaew, Shanna H. Coop, Jude F. Mitchell

**Affiliations:** Neuroscience Graduate Program, University of Rochester; Brain and Cognitive Sciences, University of Rochester

## Abstract

Primates rely on eye movements made 2-3 times every second to scan their visual environment. Each eye movement induces substantial retinal motion, yet observers readily suppress it to stitch together the percept of a stable world. Neurons in the middle temporal (MT) and medial superior temporal (MST) areas respond to motion, but also suppress that response during eye movements (Bremmer et al., 2009; Leopold & Logothetis, 1998; Thiele et al., 2002). This saccadic suppression could reflect the influence of corollary feedback from eye movement related brain structures (Berman et al., 2017). However, visual signals that detect wide-field motion characteristic of a saccade could also contribute to suppression, akin to visual masking (Idrees et al., 2020). We sought to disentangle the relative contributions of visual signals and corollary discharge in driving saccadic suppression in areas MT and MTC in marmoset monkeys. Marmosets freely viewed natural images, blank backgrounds, or externally moving images that simulated saccades, and also made saccades in complete darkness. We found a diversity of saccadic modulation across the population. Some neurons showed short latency excitatory responses to saccades while others showed suppression. Still others showed a biphasic response that began with suppression followed by an excitatory rebound. Neurons with early responses were frequently tuned for the direction of saccades on natural images and retained that same tuning for saccades on blank screens or in complete darkness, thus supporting a role for corollary feedback. However, these neurons also responded with similar directional tuning for retinal motion created by simulating saccades while the eyes were fixed, and thus also appear to integrate visual cues for saccadic motion. More so, these units were biased towards having narrow spike waveforms consistent with putative inhibitory cells, which drive suppression observed in the rest of the population. We also found a subset of neurons that responded robustly to simulated saccadic motion but had negligible response to motion from real saccades, such that they discounted saccade induced motion. Those neurons were biased to have broader spike waveforms, consistent with being putative excitatory projection neurons and could support our percept of stability during eye movements.

## Introduction

The visual world is far too large to obtain all the information in a single glance. Instead, humans and other primates use rapid eye movements, called saccades, to direct detailed visual processing toward areas of interest. Saccades are made frequently, typically 2-4 per second, and with velocities as high as 700 degrees per second, that are brief in duration (∼30-50ms), to efficiently process a scene (Baloh et al., 1975; Fischer & Weber, 1993; Wolfe et al., 2011).

However, due to the visual motion induced by each movement, saccades repeatedly create perceptual discontinuities and spatial instabilities. To perceive a stable and constant world, the visual system filters out this self-made motion, resulting in decreased visual sensitivity during saccade flight (Volkmann, 1986; Zuber et al., 1966). This phenomenon is called saccadic suppression.

Two proposed mechanisms to mediate saccadic suppression involve feedforward and feedback pathways. In the feedforward view, large changes to visual input seen during saccades, such as wide field motion and transients in contrast, trigger suppression (Castet et al., 2002; Castet & Masson, 2000). Researchers have shown that saccade-induced changes to firing rate can cause suppression in visual sensitivity as early as the retina (Idrees et al., 2020). Alternatively, in the feedback view, copies of motor planning signals (corollary discharge) are sent to visual processing areas to trigger suppression (Duffy & Lombroso, 1968; Holt, 1903; Matin, 1974; Zuber et al., 1966). Motor planning areas such as the Frontal Eye Fields (FEF), and the Superior Colliculus (SC) are candidates for triggering oculomotor feedback (Chen & Hafed, 2017; Pierrot-Deseilligny et al., 1995; Zanos et al., 2016; Krock and Moore, 2016). Both area FEF and SC have connections with visual processing areas that could mediate suppressive signaling, either directly in the case of FEF (Anderson et al., 2011) or indirectly through the pulvinar in the case of SC (Berman & Wurtz, 2011; Miura & Scanziani, 2022).

More recently, evidence in area V1 has shown that single unit responses after saccades vary in their timing and sign of modulation giving rise to complex dynamics across the neural population (Parker et al., 2023). Following the suppressive period, neural activity exhibits a post-saccadic (or re-fixation) enhancement which peaks 50-150ms after saccade onset. Diversity in the degree of suppression and the timing of rebound were correlated with the spatial frequency tuning of neurons in V1, with neurons tuned to higher spatial frequencies responding at slower latency. It was proposed that this organization in post-saccadic responses can contribute to a coarse-to-fine (low to high spatial frequency) processing strategy.

The relative contributions of visual saccade detection and corollary feedback signals in driving suppression, and how they relate to the subsequent population dynamics after saccades, are yet to be determined. Area MT and MST provides a unique opportunity to study population dynamics around saccadic suppression because they specialize in motion processing and thus would particularly need to suppress it during eye movements to support perceptual stability (Thiele et al., 2002; Chukoskie & Movshon, 2009). In particular, in the marmoset monkey both of these areas lie at the cortical surface allowing recordings from neural populations as well as for making laminar distinctions in recorded neurons to determine how diverse response properties are distributed in the population. Therefore, we investigated the diversity of saccade-related modulation in areas MT and MTC (the MST-lateral/V4t equivalent in macaques) and tested to what extent extra-retinal or visual signals contribute to saccadic suppression.

## Methods

### Subjects

All experimental procedures followed the National Institutes of Health Guide for the Care and Use of Laboratory Animals. Protocols for experimental and behavioral procedures were approved by the University of Rochester Institutional Animal Care and Use Committee. Three adult common marmosets (Callithrix jacchus), Marmoset M (male), Marmoset S (female), and Marmoset A (male) were used for neurophysiological recording experiments to measure changes in neural activity during saccades. Subjects were housed at the University of Rochester with a circadian cycle of 12-hour light and 12-hour dark.

All subjects were surgically implanted with head caps to stabilize them for head-fixed eye tracking and neural recordings as described previously (Nummela et al., 2017; Coop et al., 2024; Bucklaew et al., 2026). Marmosets were then trained to sit in a primate chair under head restraint while facing a computer screen with vision corrected by a lens (Nummela et al., 2017).

### Eye Tracking and Stimulus Presentation

Eye movements were tracked with an infrared eye tracker (EyeLink 1000, SR Research Ltd) with a precision of 0.10 visual degrees (Yates et al., 2023). Eye position was first calibrated at the start of each behavior session using a Gaussian windowed face detection task described previously (Mitchell et al., 2014; Nummela et al., 2017). The eye calibration was then fine-tuned using a fixation task with a small point (0.3 dva radius) at the screen center (Mitchell et al., 2014). Eye position data were collected during the entire recording session and smoothed offline as described in a previous study (Coop et al., 2024). Briefly, saccadic eye movements were detected offline using an automatic procedure that detected deviations in 2-D eye velocity space and marked by where the 2D velocity exceeded the median velocity by 10 SD for at least 15ms (Engbert & Mergenthaler, 2006; Kwon et al., 2019). Saccade onset and offset were determined by the first and last time the 2-D velocity crossed the median velocity threshold. Epochs with eye blinks (detected by absence of pupil size) were removed from analysis.

Stimuli were generated via Psychophysics ToolBox (Brainard, 1997; Pelli, 1997) in Matlab and presented on a gamma-corrected display (BenQ X2411z LED monitor, resolution: 1,920 x 1,080 p, refresh rate: 100Hz, gamma correction: 2.2) that had a dynamic luminance range from 0.5 to 230 cd/m2 at a distance of 57 cm in a dark room. Task events and neural responses are recorded using a DATAPixx I/O box (VPixx Technologies) for temporal registration. The MATLAB code is available online (https://github.com/jcbyts/MarmoV5) and was used in previous studies (Parker et al., 2023; Yates et al., 2023).

### Stimuli

We examined the neural firing responses time-locked to saccade onset across various stimulus presentations. Stimuli were either a natural image, grey screen, moving random dot fields, or a simulated saccade that translated a natural image on the screen with the velocity profile of a typical saccade. In the natural image condition, a set of randomly selected natural scenes were presented for 10ms at a time while the subject was allowed to freely view. In the grey screen condition, we utilized a full-field luminance stimuli that flashed for 400 ms in white (230 cd/m^2^) and returned to gray (115 cd/m^2^) for 1200 ms while subjects freely viewed. The full-field luminance changes were also used to analyze the current source density (CSD) which is described later, as is the simulated saccade condition.

Moving random dot field stimuli were used to ascertain the visual latency of each neuron. Visual receptive fields were first mapped using full field moving dot stimuli as described previously (Coop et al., 2024; Yates et al., 2023). We then showed marmosets stimuli consisting of a random dot field placed in their receptive fields while they performed a fixation task to control eye position. The direction of motion in the dot field was independently varied across trials sampling from 16 directions around the circle. Each aperture contained a field of black dots (each dot 0.15 dva diameter with a density of 2.54 dots per visual degree squared) which moved at 15 degrees/s in one direction (100% coherent) with 50ms limited lifetimes and asynchronous updating to new locations. The radius of the target aperture was set to half of the receptive field eccentricity and dot contrast was Gaussian windowed from black at the center (0.5 cd/m2) to the background gray on the edges using a Guassian sigma 1/6th of the aperture’s diameter.

To characterize single unit tuning to orientation and spatial frequency, full-field Hartley Grating Stimuli were shown to map neuron’s tuning preferences as described previously (Yates et al., 2023). During these trials, a spatial frequency between 0.5 -16 cyc/deg (in 6 equally spaced increments in log spatial frequency) and an orientation between 0 – 360° (in 12° increments) was drawn for 20 ms with random sampling of the stimuli between frames. To encourage continued viewing during flashed grating stimuli, a marmoset face (1 dva) was randomly positioned and superimposed in front of the flashing grating stimuli and the marmoset received juice for finding the face (Yates et al., 2023).

In the simulated saccade condition, a 1600×700 pixel section of the natural image (same set as used previously for foraging natural images) was moved at randomly selected times during free viewing. Each simulated saccade occurred with a random inter-saccade interval ranging from 300-700 ms (uniform distribution) and in a random direction (0-360 degrees) and amplitude (1-8 visual degress). The simulated saccade had a duration of 40ms with a Gaussian shaped velocity profile approximated over 5 points, with the peak velocity at the center point of typically 200 dva/sec or more. Each trial lasted for 10 seconds before a 2-second inter-trial interval.

### Dark Condition

In the dark condition, all monitors and lights were turned off to approximate complete darkness. The subject was draped with a blackout curtain to limit outside light from entering and the infrared light source for eye tracking was placed laterally relative to the eye, removing any IR reflections from the mirror in front of the animal. An infrared eye tracker was used (EyeLink 1000, SR Research Ltd) with pupil thresholding turned off to allow for tracking with pupil dilation in the dark. This condition was performed at the end of a recording session to prevent calibration changes in other tasks.

### Recording Parameters

After an initial head implantation surgery and chamber placement using stereotaxic coordinates for MT (Paxinos, 2012), a second surgery was performed to create a craniotomy centrally within the chamber following procedures describe previously (Bucklaew et al., 2023). Craniotomies were sealed with a 0.5-1 mm layer of silastic gel (Kwik-Sil, World Precision Instruments) to protect the brain from infection and reduce granulation tissue growth. Laminar recordings were made with multichannel linear silicon arrays that penetrated the protective layer of silastic allowing access to the underlying brain. The linear arrays had two shanks spaced 200 microns apart, each with 32 electrode channels at 35-micron inter-contact spacing (NeuroNexus).

All neurophysiology data was amplified and digitized at 30KHz with Intan Head-stages (Intan) using the Open Ephys GUI. The wideband signal was high-pass filtered by the head-stage at 0.1hz. Data was then offline spike sorted using Kilosort2 and manually labeled using the “phy” GUI. Units with physiologically implausible waveforms were classified as noise. Clusters that could not be fully separated from other clusters or the noise floor, or that exhibited more than 1% inter-spike interval violations under 1 ms, were counted as multi-unit activity.

### Receptive Field Mapping

Spatial receptive fields were estimated from neuronal responses to a wide field stimulus of moving dots as reported previously (Coop et al., 2024; Yates et al., 2023). Marmosets freely viewed a full-field display that consisted of white dots (230 cd/m2,1 dva diameter) that appeared against a gray background (115 cd/m2) spanning ±20 dva on the horizontal and ±15 dva on the vertical of the display. Each dot moved at 15° per second for 50ms before being replotted to a new location with a new motion selected at random from 1 of 16 motion directions sampled around the circle. Eye position was corrected offline to determine the receptive field properties as described previously (Yates et al., 2023). In brief, the firing rate was time-locked relative to the onset of dots across a grid of retinal spatial locations in 10ms spike counting bins. Those grid locations exhibiting firing responses significantly above the pre-stimulus baseline firing (from −100 to 0ms lag, p < 0.001) were labeled as significant. A smoothed 2D contour was then computed to circumscribe the peak of the RF at 80% of the maximum height relative to the baseline at the peak temporal lag.

### Current Source Density Analysis

In each recording we also estimated the laminar depth of electrodes by applying the Current Source Density analysis (Mitzdorf, 1985). Previous work analyzing this current source/sink in macaque V1 and MT showed that the input layer exhibits a short latency current sink followed by a reversal to a current source in response to a full-field luminance change (Schroeder et al., 1998). This method has been used to mark the bottom of the input layer, after which the top of the input layer can be estimated based on the average width from anatomical studies, estimated at 300 microns for marmoset MT (Bourne et al., 2007).

To obtain CSD estimates in our data, we showed marmosets two visual stimuli during each recording session. Similar to what is commonly used in V1, the first stimulus was 400 ms full-field luminance flashes (from 115 cd/m2 background to 230 cd/m2), presented every 1600 ms. Additionally, considering both areas being recorded have mainly been shown to be involved in motion processing, we showed marmosets full-field stimuli consisting of 640 dots (0.2 dva diameter) randomly distributed across screen locations which were held stationary for 400 ms before making full-field coherent motion in 1 of 16 random directions at 15 dva/s for 400 ms, and then repeated for 10s each time testing a different sampled motion.

LFPs were first obtained from raw neurophysiological data by low-pass filtering (300Hz, 1st order Butterworth filter) and down-sampling to 1kHz. Then, any artifacts 10 standard deviations above the median were removed along with line noise at 60, 120, and 180 Hz. To process LFP signals during CSD stimuli offline, we approximated the second spatial derivative following Mitzdorf (1985). Afterward, we smoothed the average CSD using a Gaussian filter across electrode depth (σ = 150 μm). To identify the reversal depth of the CSD in MT/MTC, we visually searched in the range of 30-50ms after stimulus onset (the visual latency of the area) for the sink location. We then selected the point of reversal between the current source and sink as the reversal location (**Supplemental Figure 2**). If we were unable to visually determine sink/source reversal in the accompanying CSD, we used reversal depth from the adjacent electrode shank or marked the depth as inconclusive if neither shank showed a clear CSD. Reversal depth identified from these analyses was set as 0 µm in all subsequent analyses to mark the bottom of input layer.

### Analysis of Neural Responses and Inclusion Criteria

A total of 1871 single units were identified. To be included in subsequent analysis, neurons were required to have: 1) a baseline firing rate of above 0.1sp/s on both the natural image and blank screen conditions, 2) a significant visual response to random dot fields shown in the neuron’s RF, and 3) a significant deviation (either excitation or suppression) of firing rate from baseline within 200ms of the onset of saccades in free viewing natural images. The peak or trough was compared to the baseline rate in the 100 ms preceding saccade onset and accepted if more than two standard deviations away. A total of 1267 units met these criteria and were included in subsequent analyses.

### Direction Selectivity Index (D.S.I.)

To assess the motion selectivity of neurons we computed a direction-selective index (D.S.I) from spike counts taken between 50 to 150ms after the random dot motion stimulus onset as

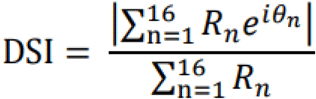

where R_n_ represents the mean spike count in response to a motion direction.

A similar method was used to calculate the direction selectivity of saccades. Saccade directions (which were randomly spread across 360 degrees) were first sorted into 12 bins (representing 30 degrees each). Then we computed the mean response for each bin, taken either in a 50ms counting bin centered on the post-saccadic peak or trough of the response. The resulting DSI for saccades and dot motion stimuli was compared against a distribution of DSI values generated by bootstrapping by randomly permuting the directions of saccades (1000 samples) and computing the mean shuffle-corrected DSI and its standard deviation. Units with DSI values having p values above 0.01 compared to the shuffle corrected distribution were considered significant.

### Simulated Saccade AI Score

To assess the relative selectivity for externally generated motion to saccade generated motion, we computed a simulated saccade AI score (SSAI) from spike counts taken 40ms to 160ms after a real saccade or a simulated saccade as

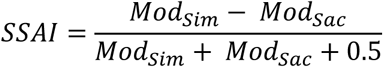

where *Mod* represents the change in baseline normalized firing rate averaged over the 40-160 ms interval for either real or simulated saccades. To avoid amplification of noise among neurons with very modest changes in rate during saccades, we add the value of 0.5 in the denominator thus emphasizing units with greater than 50% modulation in at least one of the conditions. Neurons were then identified as External Motion Units if the SSAI was greater than 2/3.

### PCA Cluster Analysis

To cluster units based on similar saccadic response profiles, we first performed a PCA on the baseline normalized saccadic responses under natural image and blank screen conditions in the period from -100 to 200ms from saccadic onset. Units included in the analysis (n = 1225) were required to have a minimum firing rate of 0.1sp/s and a significant saccadic response that differed from basline (100 ms before saccade onset) to be included in the analysis. A small proportion of units was also excluded from this analysis if they showed saccade modulation (peak or trough) that was less than 25% of baseline (n = 42, 3.3%). We then used the first four components of a PCA analysis, which cumulatively explained 95% of the variance, to perform a K-means clustering following a previous study (Parker et al., 2023). This resulted in three clusters of unit response profiles to best explain the data variance with minimal number of clusters (**Supplemental Figure 1**).

## Results

### Neural Modulation During Saccades in Areas MT/MTC

To investigate saccade-related modulations of neural firing in areas MT/MTC, marmosets freely-viewed natural images or blank screens displayed 10 seconds at a time (**Figure 1A, top two panels**). While subjects made spontaneous saccades, we simultaneously recorded neural activity in areas MT and MTC using silicon linear array probes. We identified a total of 1871 well-isolated single units in three animals (M = 344, S = 557, A= 970). We also tested their response to visual motion by presenting random dot motion stimuli inside their receptive fields while the marmoset was fixating (**Figure 1A, bottom panel)**. For initial analysis, neurons were only included if they exhibited a significant saccade-related modulation, either with a peak and/or a dip in firing after saccades, that deviated by more than 2 standard deviations relative to baseline firing in the 100 ms epoch before the saccade. Collectively from three animals, this resulted in a total of 1267 neurons (M = 232, S = 366, A = 669).

**Figure 1:**
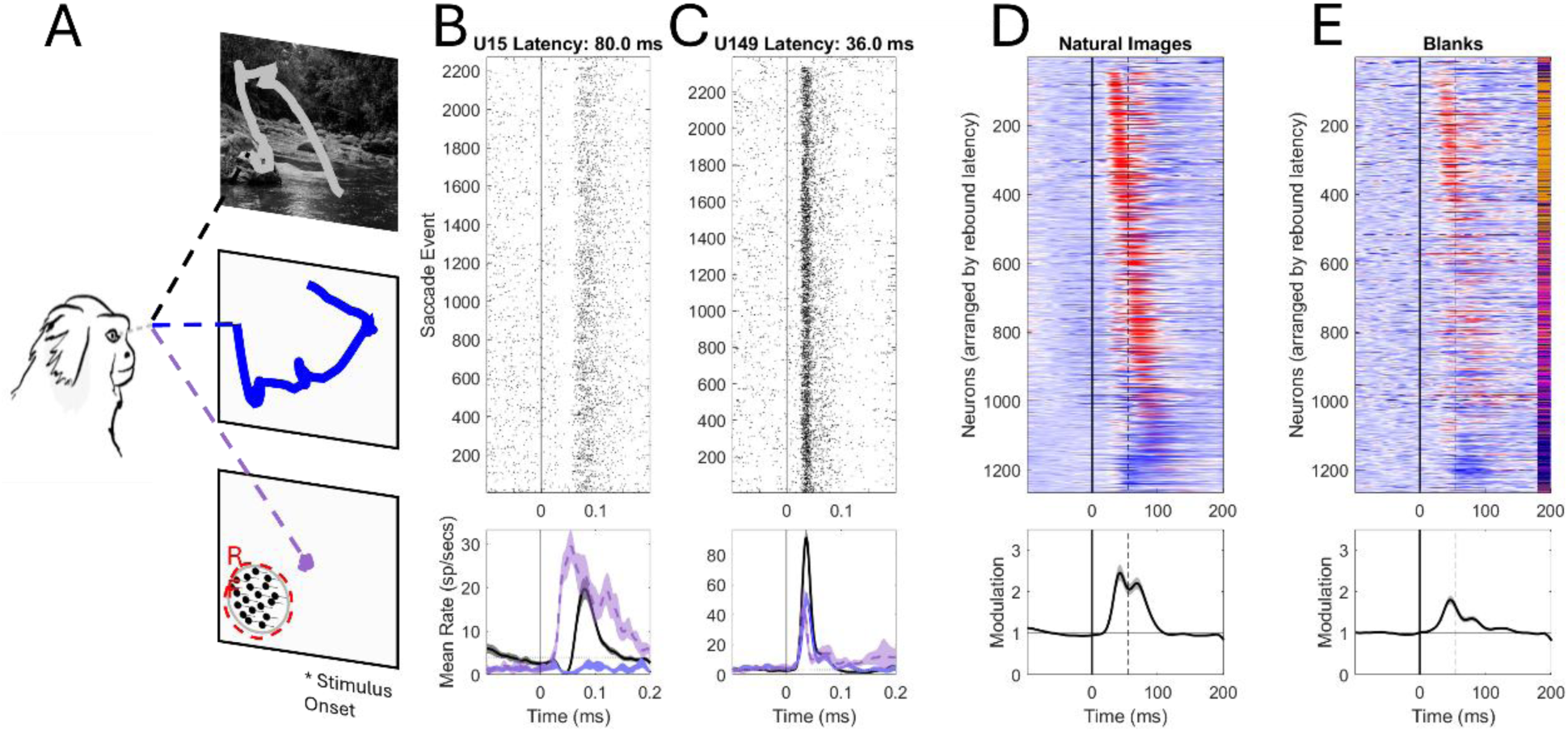
Neural Responses during Saccades on Natural Images and Blank Screens. A) Marmosets freely viewed either natural images or blank screens for 10 seconds at a time or held fixation while a motion dot field was presented in the neuron’s receptive field B-C: Top) Example unit PSTH for the natural image condition (black) aligned to saccade onset at time zero B-C: Bottom) Example Unit mean normalized firing rate for natural image conditions (black, aligned to saccade onset), blank screen conditions (blue, aligned to saccade onset), and visual response to motion dot fields in the receptive field (purple, aligned to stimulus presentation). D) Natural Images PSTH of normalized saccadic modulation wherein suppression below baseline is shown in blue and excitations above baseline shown in red. E) Blank Images PSTH of normalized saccadic modulation wherein suppression below baseline is shown in blue and excitations above baseline shown in red. Identifiers on right show the cluster each unit was classified into.

One issue in measuring saccade modulation on natural images is that each saccade creates retinal motion that will drive visual responses from MT neurons. The resulting modulation is thus potentially representative both of an extra-retinal response to motor feedback from the movement, but also a visual response to retinal motion. To better dissociate the visual from extra-retinal component, we additionally had marmosets free view blank screens (**Figure 1A, middle panel**). The blank screen stimulus reduces changes in visual input during saccades because it reduces motion signals within the receptive fields. However, the blank screen is not a perfect control for saccade-induced motion because the edges of the screen are still present and could influence responses from the surround of receptive fields. As further control, in one of the three marmosets, we recorded while the marmoset made saccades in the dark.

Across our population, we saw a variety of neural responses following saccade onset (**Figure 1B-C)**. An example unit exhibited classical patterns of saccadic suppression with an initial suppression followed by a post-saccadic peak (**Figure 1B**, **black curve**). This unit instead showed a purely suppressive response when making saccades on a blank screen (**Figure 1B, blue curve**). Additionally, when a motion stimulus was presented in the example neuron’s receptive field, it showed a sharp response at a short visual latency that was aligned with the start of suppression in the natural image and blank conditions (**Figure 1B**, **purple curve**).

Another example unit, by contrast, showed an early excitatory response to saccades with no sign of suppression (**Figure 3.1C**, **black curve**). In that unit, we observed increases in firing rate at a similar latency in the blank screen condition (**Figure 1C**, **blue curve**), and this response was consistent with the response latency to visual motion onset in the receptive field (**Figure 1C, purple curve)**. These purely excitatory saccade-related responses occurred rapidly near the neuron’s visual latency and thus could reflect a visual response to the saccade-induced retinal motion. However, it is interesting that the short latency response also persisted for the blank screen condition which would support a role for corollary feedback.

Across the population we observed a continuum of saccade-related response modulations ranging from early responses lacking suppression to responses that began later, and in some cases, only showed suppression (**Figure 1D**). To visualize the responses across the population we plotted the mean normalized saccade response relative to baseline (baseline modulation = 1), with suppression shown in blue and excitation in red, and with neurons ordered on the vertical axis by the latency of their peak post-saccadic response. Under Natural Image conditions (where visual input is high), we observed a bimodal distribution of peak responses wherein some units had an early peak under 50 ms from saccade onset, and across the full population this interval showed a mean positive modulation 145% above baseline (rank-sum, p<0.001). Other neurons had a late firing rate modulation occurring after 50ms from saccade onset, and across the full population this interval showed a mean positive modulation 123% above baseline (rank-sum, p<0.001). Still other units exhibited a purely suppressive response with no excitatory rebound (**Figure 1D, bottom**).

The saccadic modulation on a blank screen produced patterns that distinguished early response units from late and suppressive units (**Figure 1E**). The modulation across the full population in the early time window (within 50ms of saccade onset) showed a population wide increase in firing rate that was driven by the subset of neurons showing an early peak response. It retained a mean positive modulation 80% above baseline with blanks (rank-sum, p<0.001). By contrast, the response across the population in the late window (after 50ms from saccade onset) was substantially diminished at only 35% above baseline (rank-sum, p<0.001). And the smaller set of units with purely suppressive responses maintained a significant suppression on blank screens. The variation in how different units respond to the blank condition suggests possible distinctions in how cells are driven by visual and extra-retinal input during saccades.

### Clustering Analysis Shows Variation of Response Types

The variance in saccade-related modulations was best explained by a set of three clusters linked in a continuous distribution (**Figure 2**). To distinguish between different response patterns, we utilized a cluster analysis applied previously in V1 data (Parker et al., 2023). Specifically, we used a K-means clustering based on principal component analysis (PCA) of the normalized peri-stimulus time histograms (PSTH) to the saccade response on natural images and blank screens. Visualizing the responses in a 3-D PCA space revealed a widespread distribution but without separate clusters, supporting a continuum of response properties that was consistent with the previous V1 study (**Figure 2A-B**). In order to break up the continuum for visualization, we submitted the data to a K-means clustering and determined the optimal number of clusters to explain the data. We varied the number of clusters fitted in the first three principal components and calculated the difference in data variance explained when adding a cluster, or the marginal gain, and then compared against the “Heuristic Threshold” of 5% gain. This resulted in an optimal value of three clusters (**Supplemental Figure 1**). The previous study had identified an optimal value of 4 clusters in V1 data. Although our MT/MTC data identified 3 clusters, the qualitative aspects of the groups identified were similar.

**Figure 2:**
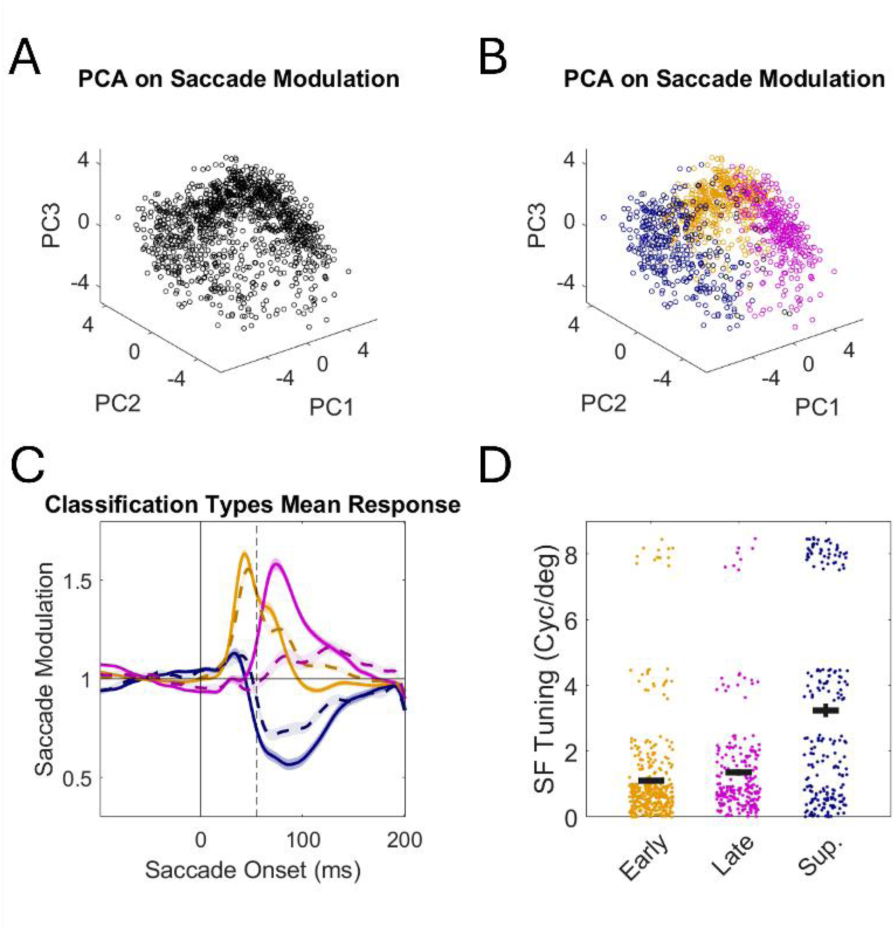
Clustering Analysis Shows Variation of Response Types. A-B) PCA and K-means clustering analysis on population responses in natural image conditions either (A) without clusters labeled or (B) with clusters color labeled. C) Mean response profiles of the three main response profile types identified by the clustering analysis D) Spatial frequency tuning of the three main response profiles identified.

The mean modulation of the three identified groups varied in response latency and degree of suppression (**Figure 2C, solid lines**). One group demonstrated an early positive response at a latency under 50 ms (n = 494, 39%). The second group also showed a positive response, but in a later interval after 50 ms (n=361, 28.5%), which we termed a late response. The final response pattern was entirely suppressive (n = 370, 29.2%). The mean modulation on blank images (**Figure 2C, dashed lines**) revealed that early response and suppressive units show roughly similar modulations for blanks, whereas the late response units showed a reduced response that was barely different from baseline (**Figure 2C, dashed lines**). Relating these responses back to the population PSTH diagram in **Figure 1E** (denoting each group by the color on the right vertical axis), we observed that the early units (labeled in orange) were responsible for the positive responses in the blank conditions, whereas late response units (labeled in purple) lacked much response, and suppressive units (labeled in blue) showed suppression in either viewing condition.

The diversity in response latency of our MT/MTC population correlated with spatial frequency preference consistent with a coarse-to-fine processing strategy found previously in V1 data (**Figure 2D**). In each recording session we estimated the spatial frequency tuning of individual MT/MTC neurons based on their response to rapidly flashed grating stimuli of varying orientation and spatial frequency ranging from 1, 2, 4, to 8 cycles per degree (see Methods).

We found that early and late responses showed preferred tuning for low spatial frequencies whereas suppressive units were tuned for higher spatial frequencies (**Figure 2D**), compatible with that seen previously in marmoset V1 (Parker et al., 2023). However, the proportions of units in the MT/MTC dataset favor early response types more so than that found in V1. The proportion of early response in MT/MTC reached 39% as compared to 19% in V1 (Parker et al., 2023). This difference may reflect underlying differences in the visual input pathways to V1 and MT. While V1 pools both from magno- and parvo-cellular inputs received from the lateral geniculate nucleus, which respectively are tuned for low and high spatial frequencies, MT is thought to primarily pool from pathways carrying magno-cellular input (Maunsell et al., 1990).

### Early Response Neurons are Preferentially Tuned to the Direction of Saccade

Since the early response units retained significant positive responses for the blank condition, we tested if they might be preferentially driven by extra-retinal inputs. For example, they might be tuned for motor features of the saccade, like its direction, regardless if it occurred for the natural image condition, with strong image motion, or the blank screen conditions, with minimal image motion. We analyzed each unit’s firing rate in the post-saccadic peak or trough as a function of saccade direction (in bins of 45 degrees over 360 degrees, 30 ms counting window centered on peak or trough). To assess the strength of directional tuning we computed a direction selectivity index (DSI), which ranges from 0 for equal responses to all directions up to 1 for responses to a single direction (see Methods). An example early response neuron exhibited a strong peak that following saccades (**Figure 3A**) which exhibited strong tuning for the direction of saccade. For saccades on natural images, it preferred upward saccades (**Figure 3B, polar plot)**, with highly significant directional tuning (DSI = 0.475, p<0.001). Although the mean firing rate was substantially diminished in the blank screen conditions, the response of this unit retained similar directional tuning for upward saccades (**Figure 3C**), that was highly significant (DSI = 0.286, p<0.001). Other response types also showed significant tuning for the saccade direction. An example late response neuron had strong direction tuning that remained consistent between natural image and blank conditions with tuning for rightward saccades (**Figure 3D-F**) and remained significant in both conditions (natural: DSI = 0.590, p<0.001; blank: DSI = 0.346, p<0.001). Among the units with purely suppressive responses, directional tuning for saccades was rare. Since those neurons lacked a peak, we instead measured the tuning from the trough of their saccade-related response (**Figure 3G**). A typical suppressive unit lacked saccade direction tuning for both the natural image and blank conditions (**Figure 3G-H**), reflected by smaller directional selective indices (natural: DSI = 0.079, p>0.05; blank: DSI = 0.119, p>0.05).

**Figure 3:**
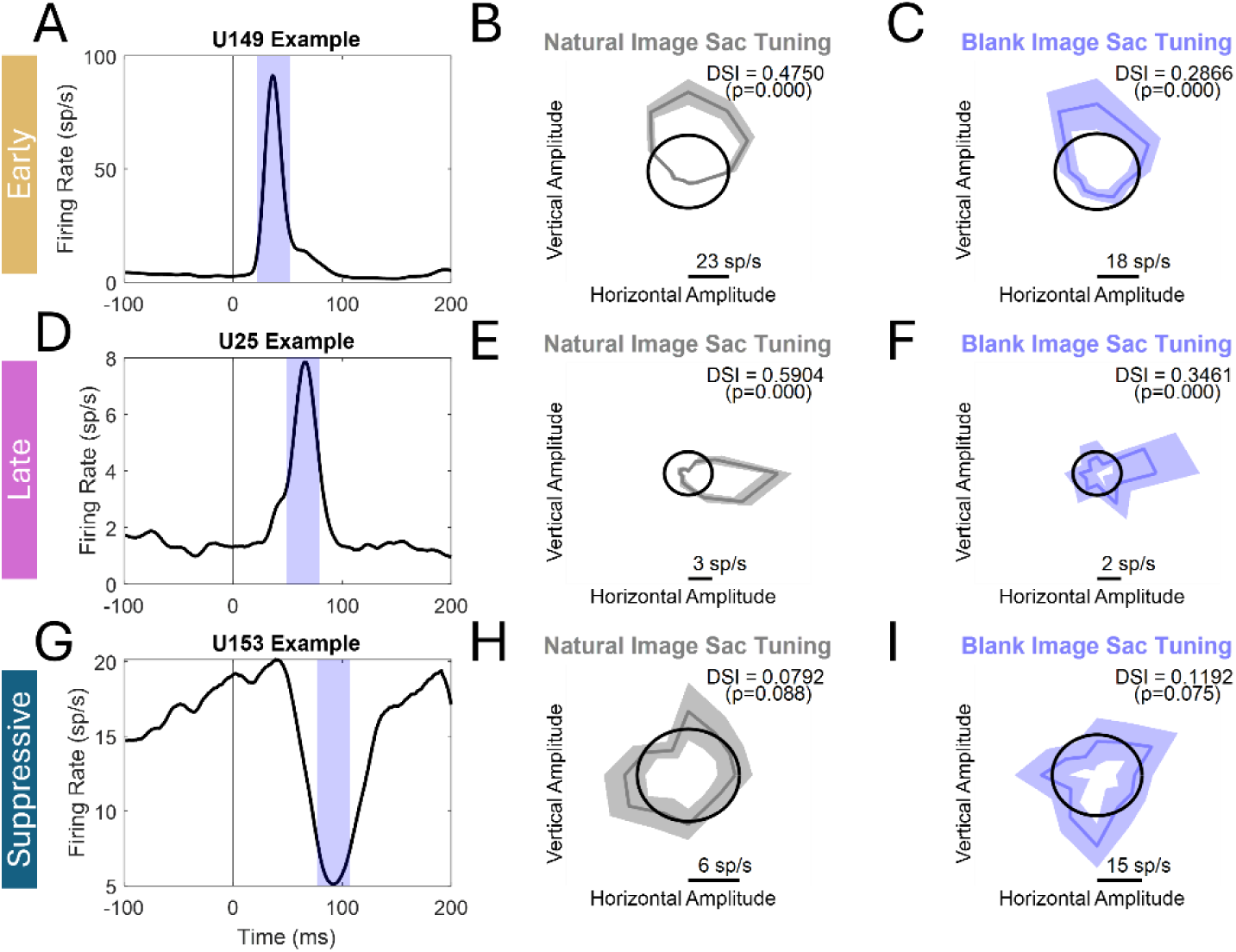
Single Unit Examples of Saccade Tuning. A,D,G) Mean firing rate time-locked to saccade onset for an example single units classified as Early(A). Late(D), or Suppressive(G). Shaded window shows identified peak or dip +/- 15ms. B,E,H) Saccade tuning direction during the Natural Image condition for an example early (B), late (E), or suppressive (H) unit. Black circles represent a uniform firing rate across saccade directions. Grey line shows the early unit’s firing rate across saccade directions. C,F,I) Saccade tuning direction during Blank Image Conditions for an early (C), late(F), or suppressive (I) unit. Black circles represent a uniform firing rate across saccade directions. Blue line shows the early unit’s firing rate across saccade directions.

Across the population we found that early response neurons showed stronger direction selectivity than neurons with either late or suppressive responses, and more so, showed a higher proportion of units with significant directional tuning in the blank screen viewing condition. The distribution of DSI values were biased towards lower values between 0.1 to 0.2 for natural images and blank screens (**Figure 4A-B**) with a significant proportion having individually significant tuning (full distribution in grey, significant labeled in color). All three response types showed some fraction of directionally tuned units in the natural image condition where visual motion might drive their selectivity (Early: 73.5%, Late: 56.5%, Suppressive: 51.1%). But even in the blank conditions, where visual motion is reduced, all response types showed more tuning than would be expected by chance at a 5% level (Early: 36.7%, Late: 24.4%, Suppressive: 20.0%). Early units had significantly stronger tuning in the natural image condition (Mean DSI: 0.167) than late (Mean DSI: 0.091, kstest, p < 0.001) or suppressive units (Mean DSI: 0.073, kstest, p < 0.001). Likewise, they had significantly stronger tuning for blank images (Mean DSI: 0.112) than suppressive units (Mean DSI: 0.048, early vs suppressive, kstest, p < 0.001) but did not differ in mean from late units (Mean DSI: 0.101, kstest, p = 0.620), though they did in proportion of tuned units (early vs late: 36.7% vs 24.4%, p < 0.001). In brief, early responses units had the strongest tuning for saccade direction.

**Figure 4:**
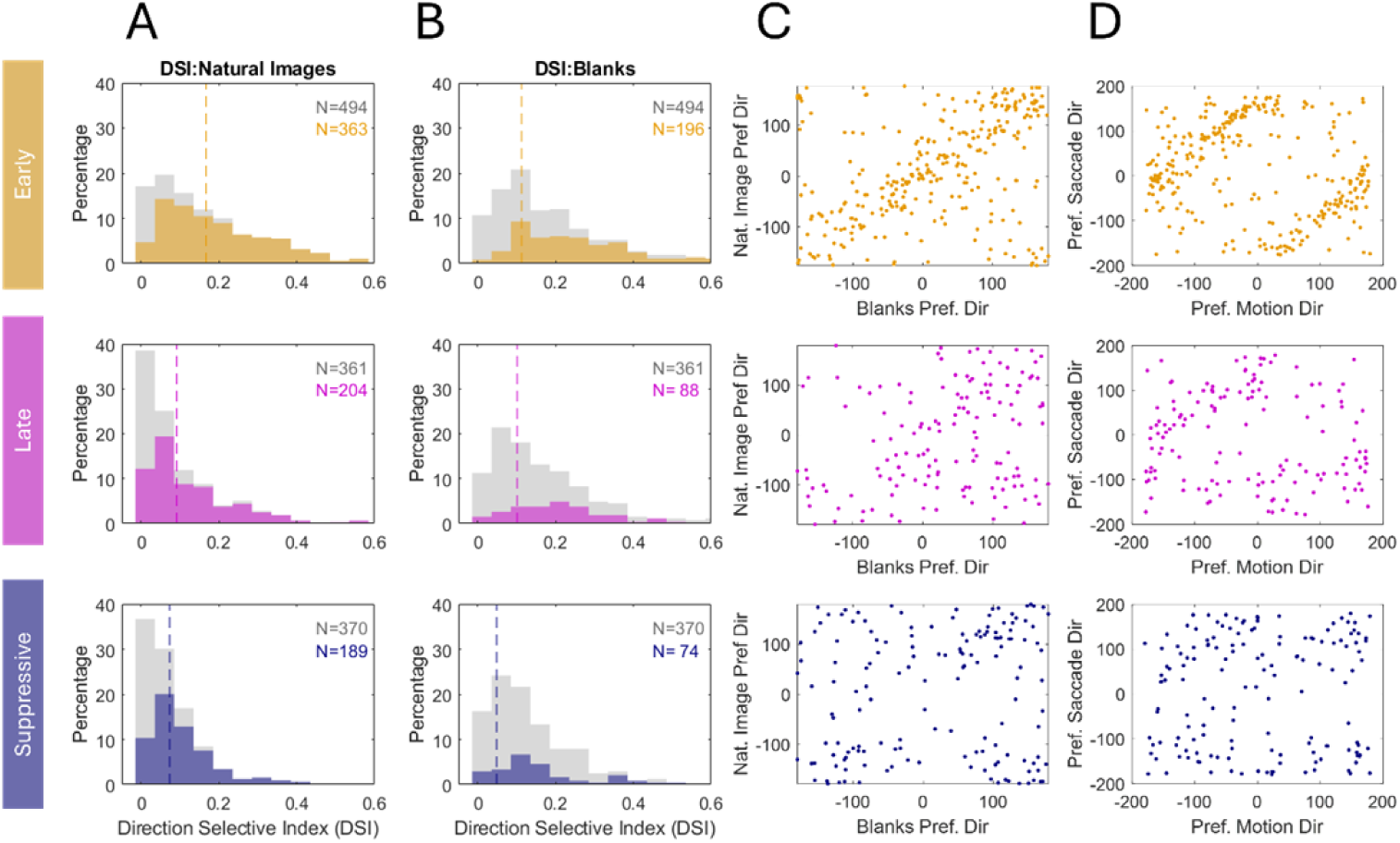
Saccade Direction and Motion Direction Tuning Show Anti-Alignment. A-B) Histograms showing the distribution of DSI strength across each cluster type for saccade tuning on (A) Natural Images or (B) blanks. The full population distribution is shown in grey, and the distribution among units with significant tuning is shown in color. Dashed vertical lines show the mean of the full distribution. C) Scatterplots showing correlation between saccade preferred direction on natural images to saccade preferred direction on blank images separated out classification type. D) Scatterplots showing correlation between saccade preferred direction and motion preferred direction separated out by classification type.

Among those neurons with significant saccade tuning there was a general alignment in their preferred direction for the natural image and blank conditions, particularly among the early response units (**Figure 4C**). Early response units showed a strong correlation between the preferred saccade direction for natural images and blank images, indicated by points falling along a line of unity in comparing preferred directions (**Figure 4C, top**). To evaluate the significance of this correlation, we used circular statistics to compute the coherence between preferred directions, a measure of correlation for a circular variable, and phase, a measure of the mean relation in radians between two sets of angles. Tuning direction in the natural image and blank conditions were significantly correlated for early units (coh = 0.555, pha = -0.083, p < 0.001), whereas for units with late responses (**Figure 4C, middle**) the correlation was much weaker (coh = 0.230, pha = 0.190, p < 0.001) as well as for suppressive units (coh = 0.236, pha = -0.387, p < 0.001) (**Figure 4C, bottom).** The correlation among early response units was nearly double that of late response (Z = 3.814, p < 0.001) or suppressive units (Z = 3.801, p < 0.001), while late response and suppressive units did not differ from each other (Z = -0.054, p = 0.9562). Thus, while all types showed some correlation in preferred tuning direction between natural image and blank conditions, early response units had the strongest correlation.

Although early response units had stronger tuning for saccade direction than other types, their tuning for motion direction was actually no different from other types. All groups showed a relatively large proportion of units with significant tuning to visual motion direction when random dot motion fields were placed in their receptive fields (early: n = 296, 81.5%, late: n = 143, 70.1%, suppressive: n = 149, 78.8%). The proportion with significant tuning for early units was greater than late response units (chi= 9.787, p=0.0018) but no different than suppressive units (chi = 3.923, p =0.0476), and neither did late vs suppressive units differ (chi = 0.583, p = 0.4453). Collectively, all of the groups show selectivity to random dot motion consistent with previous studies in areas MT and MTC of the marmoset monkey (Rosa & Elston, 1998), and this was not correlated with their type of saccadic modulation.

We observed an anti-alignment between the direction of visual motion preference and saccade direction preference, which is consistent with the saccade tuning being driven by a response to visual motion (**Figure 4D**). Specifically, a saccade made to the right will induce a leftward shift of image features on the retina, and thus have an opposite direction of tuning. An anti-aligned tuning preference thus supports a visual explanation of the response as opposed to extra-retinal corollary discharge. We found that saccade tuning was strongly anti-aligned with motion tuning for early units (coh = 0.567, pha = -2.814, p < 0.001), with weaker anti-alignment for late response units (coh = 0.365, pha = -2.630, p < 0.001), and no significant relationship for the suppressive units (coh = 0.100, pha = -1.539, p = 0.226). Consistent with an opposite alignment, the phase was nearly opposite (-2.76 and -2.81 radians) for both early and late response units where there was a significant correlation. This suggested that the saccade-related direction dependence was related to a visual response for the saccade induced retinal motion, possibly with little or no contribution from extra-retinal inputs.

### Disentangling Visual Input and Extra-retinal Input in Early Responses

To isolate possible extra-retinal contributions to saccade modulation, in a single marmoset we examined the saccade modulation in complete darkness. At the end of each recording session, all light sources were removed from the recording rig, and the infra-red diodes used for eye tracking were repositioned outside the marmoset’s field of view. The marmoset was encouraged to free view in the dark with intermittent rewards provided for finding invisible targets on the screen, while we recorded from areas MT/MTC (A=669). Units were separated into early (n= 316), late (n=77), and suppressive (n = 250) responses as described above.

The response in the dark was weak across the population, but a subset of early response units did have a significant response supporting a role for extra-retinal inputs (**Figure 5A-B**). From the 669 units that showed a significant modulation for saccades on natural images, we selected the subset (N = 199) that showed significant saccade direction tuning and looked at their dark response. Most of these units had weak modulation in the dark (**Figure 5A**), and the average modulation remained relatively flat close to baseline (**Figure 5B**). However, there was a significant increase in the mean relative to baseline in the early period before 50 ms (11% increase, rank-sum, p< 0.001) and in late period after 50ms (9% increase, rank-sum, p = 0.002). This weak response reflects that some units showed significant positive responses. A set of units with early responses, that we will term “Early Dark” units, showed highly significant responses. Averaging across all the units of that response type, we found positive response to saccades in the dark that peaked at a later time point just after 50ms (**Figure 5C**). These dark responses showed a significant saccade tuning that correlated with the natural image saccade tuning (**Figure 5D**), which was highly significant (coh = 0.630, pha= -0.070, p < 0.001). By contrast, the correlation of preferred saccade tuning for late units and suppressive units (not shown) was not significant (late units: coh = 0.354, pha = -0.400, p = 0.343; suppressive units: coh = 0.148, pha = -0.915, p = 0.588). Thus, at least some early response units did show responses in the dark that were consistent with an extra-retinal input and maintained the same direction of tuning as seen for saccades in the light, although at a slower latency than the response in light.

**Figure 5:**
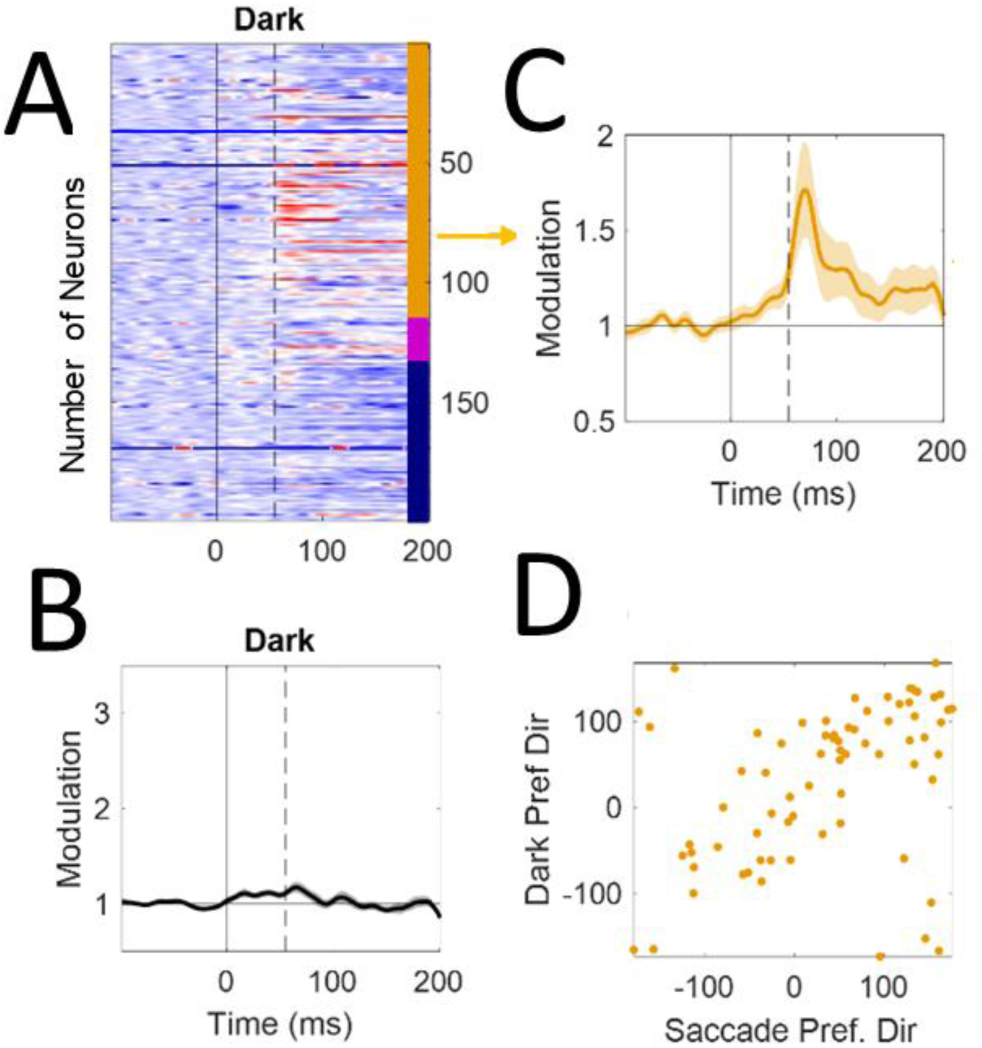
Saccadic Modulation in the Dark for Area MT/MTC is limited to a subset of neurons with early Responses. A: PSTH of modulation for all units only during the dark condition wherein suppression below baseline is shown in blue and excitation above baseline is shown in red. B: Mean modulation response of neurons during dark condition time locked to saccade onset. C: Mean modulation response of early units only during the dark condition time locked to saccade onset. D) Scatter plot showing correlation between dark preferred direction and saccade preferred direction on natural images for early units.

### Distinguishing Between Real and Simulated Saccades

In a previous study of area MT and MST (Thiele et al., 2002), researchers found a subpopulation of neurons that were selectively silenced during saccades but responded well to similar external image motion from shifts that simulate saccades. We tested if units in our population might share that property, and if so, how they would be distributed across the response types defined previously. We had a single marmoset freely view a natural image while at randomly spaced intervals we simulated saccades by moving the natural image at a typical speed and duration for marmoset saccades. The saccade duration was fixed at 40 ms with a Gaussian shaped velocity profile. The saccade direction was sampled at random with amplitudes ranging uniformly from 1-8 degrees.

The modulation for simulated saccades mimicked what we have seen previously under saccadic conditions, and in fact, even produced an increased response relative to real saccades (**Figure 6A**). On average, simulated saccades produced a peak response in the 30-80 ms post-saccadic interval that was 55% times greater than that of real saccades (Sim vs Real: 2.86+/-0.11 vs 1.84+/-0.06, p<0.001). When the population was split into early, late, and suppressive groups (**Figure 6B-D**), all groups showed some suppression for real saccades (dashed lines) relative to simulated saccadic motion (solid lines), and this reduction was significant for each group (Sim vs. Real, Early: 4.67 vs. 3.15, p<0.001; Late: 2.72 vs 2.19, p=0.010; Suppressive: 1.13 vs 0.83, p<0.001). Among early response units (**Figure 6B**), the evoked response to simulated saccades was strongly peaked (4.67+/-0.22) , resembling that of real saccades (average 3.15+/-0.16).

**Figure 6:**
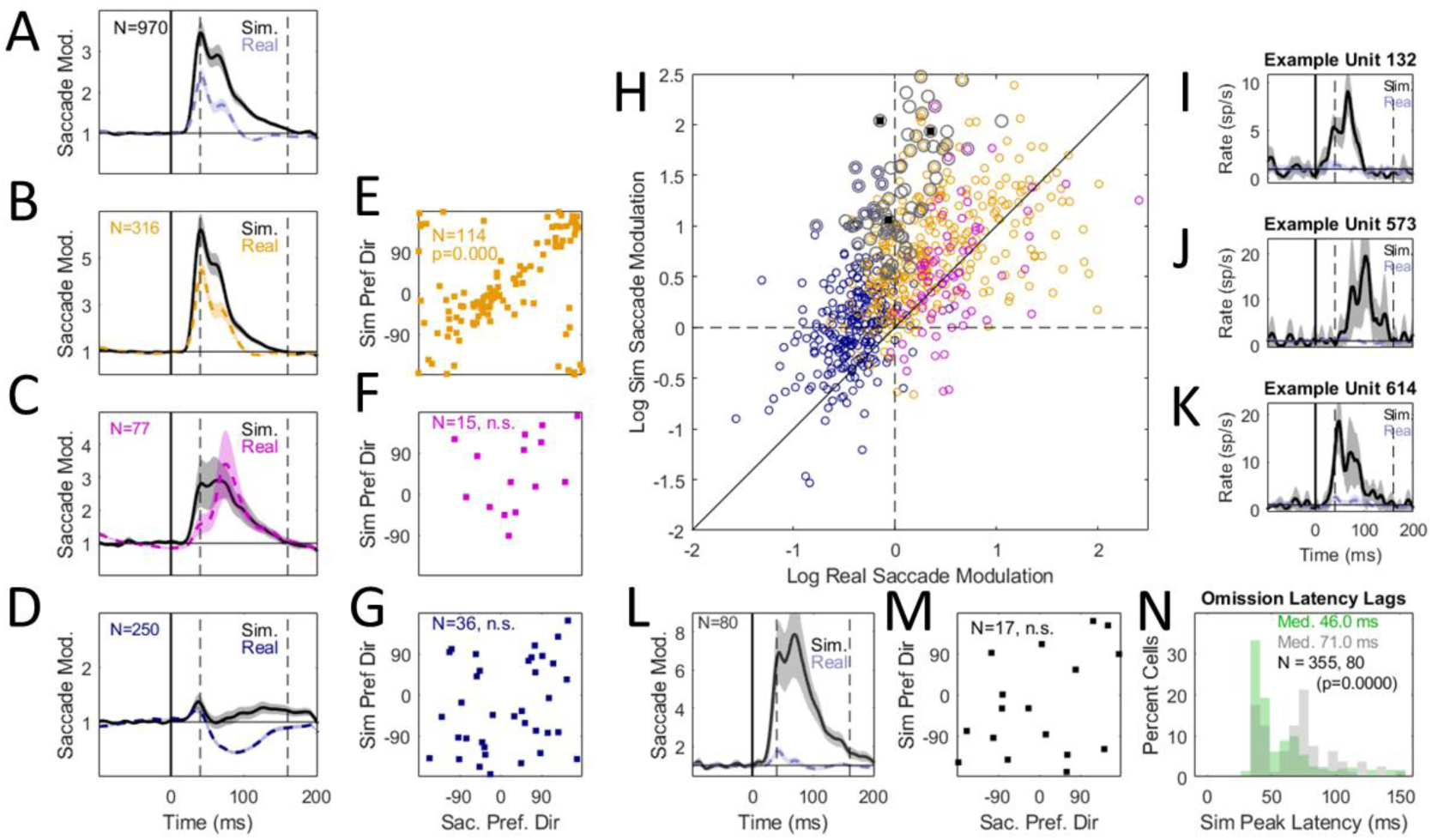
Subpopulation Responds to Simulated but not Real Saccades. A) The population mean normalized saccade modulation on natural images was reduced for real (blue, dashed line) as compared to simulated (black, solid line) saccades. B-D) Mean normalized modulation for simulated vs. real saccades broken out for (E) early, (F) late, and (G) suppressive units. E-G). The correlation of the saccade direction preference for real vs. simulated saccades was significant for early units (E) but not late (F) or suppressive units (G). Scatter plots only include units with a significant direction preference for both real and simulated saccades (Early: N = 114 of 31c; Late: N = 15 of 117; Suppressive: N = 3c of 250). H) Scatter plot showing peak modulation for simulated vs real saccades for early (yellow), late (pink), and suppressive (blue) units. Units identified with strong preference for simulated but not real saccades (see methods) are shown with grey circles. I-K) Single unit examples with strong responses to simulated but not real saccades. L) The mean normalized saccade modulation among the subset of “saccadic omission” units identified with strong responses to simulated but not real saccades. M) Unit with saccadic omission lacked a significant correlation between saccade direction preference for simulated and real saccades. N) Histogram showing peak modulation latency for simulated saccades for the saccadic omission units (gray) compared to the rest of the population showing positive responses to simulated saccades (responses).

This suggests that much of the saccadic modulation in early units might be driven by image motion. Consistent with that, we found that those early response units which had significant directional tuning for simulated and real saccades (N = 114 of 336) had a strong alignment in their directional preferences (**Figure 6E**) which was highly significant (coh = 0.74, pha = -0.12, p < 0.001). By contrast, those late and suppressive units that had significant direction tuning for simulated saccades was smaller in proportion (late: N = 15 of 77; suppressive: N = 36 of 250) and lacked alignment in directional preference with real saccades (**Figure 6F,G**), which was not significant for either group (late: coh = 0.25, pha = 0.32, p = 0.396; suppressive: coh = 0.17, pha = 0.82, p = 0.349). Thus, the visual drive influencing saccadic modulation is particularly strong among early response units, although all groups showed stronger responses to simulated than real saccades.

Across the total population we found a wide variation in the responses to real and simulated saccades which included a subset of units that omitted the response to motion from real but not simulated saccades. The mean response for real versus simulated saccades in the post-saccadic interval (30-80 ms) varied from positive and peaked to suppressive, but otherwise showed a continuum generally with stronger activity for simulated saccades (**Figure 6H**). In that distribution, we noted that several neurons exhibited a robust response to simulated saccades but almost no response to real saccades, as illustrated for three example units (**Figure 6I-K**). As those units suppressed a motion response when it is caused by a real saccade, they exhibit “saccadic omission” (i.e, they responded only to externally induced retinal motion and not self-induced motion). To identify these “saccadic omission” units, we defined a Simulated Saccade Index (SSI) as the baseline normalized response to Simulated Saccades minus that to Real Saccades, normalized by the sum of both plus a small constant (see Methods). Neurons strongly selective to simulated but not real saccades will show a value near 1, while those that do not distinguish between them will have values near 0, or negative values if they respond preferentially to real saccades. In the population we identified 80 units with strong SSI scores (SSI > 0.66) that we labeled as omission units (**Figure 6H, gray circles**).

Omission units were not distinct to any of the response types that we previously marked using cluster analysis, distributing roughly equally at the intersection between early and suppressive response types (**Figure 6H**). We found that most of the omission units were not even included in our original clustering analysis because they did not exhibit a significant response to real saccades on natural images or blank screens (included: 40%, 32 of 80; excluded: 60%, 48 of 80). Of those units that were included in the cluster analysis, most were in the early and suppressive labeled categories (Early: 59.4%, Late: 6.2%, Suppressive: 34.4%). If we averaged the modulation across the omission units (**Figure 6L**), we found the response to the real saccade was nearly eliminated relative to simulated saccades (Sim: 6.6+/-0.69, Real: 1.37+/-0.07, p<0.001). Omission units generally lacked directional tuning for real saccades (N = 18 of 80) and when tuned, were poorly aligned in directional preference between real and simulated saccades (**Figure 6M**) and not significantly correlated (coh = 0.28, pha = -0.74, p = 0.260).

Thus, unlike early units, they do not appear to code for saccade direction. Another feature that distinguished omission units was that they had slower visual response latencies to simulated saccades (**Figure 6N**), which was significant (Latency_Sub_= 71.0ms, Latency_pop_ = 46.0ms, kstest, p<0.001). Thus, while early response units could serve to detect saccades, either using visual or extra-retinal cues, omission units responded at a slower latency to retinal motion and were able to discount it when caused by a real saccade.

### Narrow and Broad Spiking Cells Differ in Laminar Distribution and Timing of Saccadic Modulation

Recording from laminar probes allowed us to investigate how saccadic modulation varied across cortical layers and cell types based on their extracellular spike waveforms. We used CSD analysis (see methods, **Supplemental Figure 2**) to identify the laminar location of recorded neurons and assign them to superficial, input, and deep layers following previous approaches used in macaque V1 and MT (Schroeder et al., 1998). We further used the spike waveform duration to split our population into putative excitatory and putative inhibitory neuronal subtypes across the different cortical layers. Previous studies suggest that narrower action potentials are on average associated with fast-spiking PV-expressing interneurons (Connors & Gutnick, 1990; Mitchell et al., 2007), while broader spikes should largely be excitatory cells.

We observed a bimodal distribution that favored broader action potential durations in the superficial and deep layers but favored narrow spikes in the input layer (**Figure 7A**). The distribution of spike waveform durations was significantly bimodal when pooled across all layers (Hartigan’s Dip Test: p=0.042), and significant in input and deep layers while approaching significance in the superficial layer (Hartigan’s Dip Test: superficial, p = 0.058; input, p=0.049; deep, p = 0.047). We separated the population into narrow (<400 us) and broad (>467 us) spike waveforms, leaving those units between the thresholds unclassified. By default, we would have anticipated nearly 80% of the population to be broad spiking based on the relative frequencies of excitatory compared inhibitory cells in cortex (Hendry et al., 1987). However, the input layer contained a large proportion of narrow spiking units far outside this range and which differed as compared to the superficial or deep layers (percent narrow: input, 44.5%; superficial, 24.5%; deep, 27.7%; kstest on spike duration distributions: input vs. superficial, p<0.001; input vs. deep, p<0.001; superficial vs. deep, p = 0.560). The proportion of units tuned for saccade direction also differed by layers. The input and deep layers included higher proportions of units with significant tuning for saccade direction in the blank image condition (marked in black in the histograms), which was a significant difference between the input and deep layers as compared to the superficial layers (proportion significant: input, 41.3%; superficial, 28.5%; deep, 43.9%; test on difference in proportions: input vs. superficial, p < 0.001 ; input vs. deep, p = 0.340, superficial vs. deep, p < 0.001). Thus, there was greater tuning for saccade direction in the input/deep layers as compared to superficial layers.

**Figure 7:**
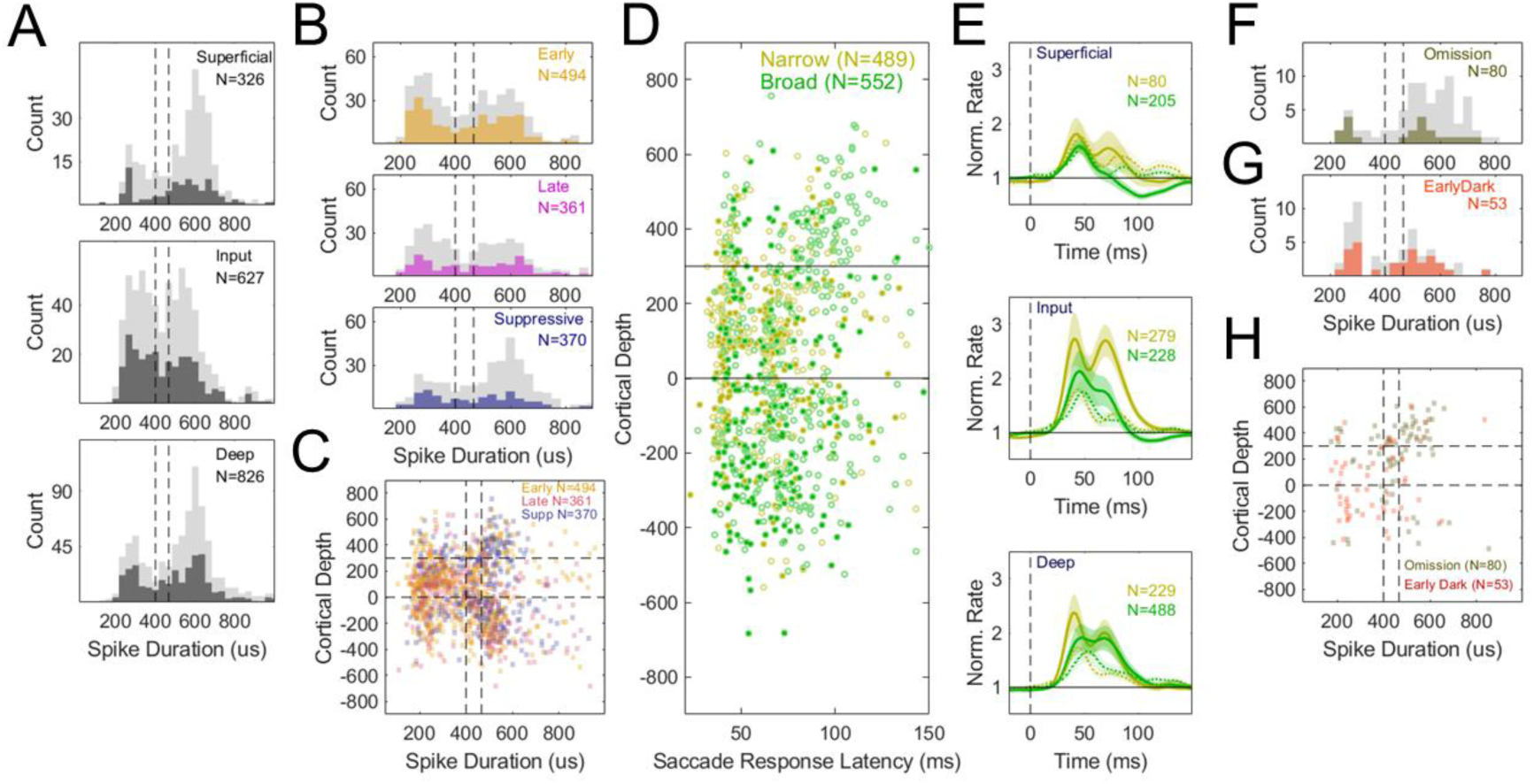
Saccadic Modulation varies across Waveform Duration and Laminar Depth. A) Histograms showing distribution of spike waveform durations for Superficial (Top), Input (Middle) and Deep (Bottom) Layers. Light grey represents all neurons while dark grey marks units with significant saccade tuning to blank images. B) Histograms showing distribution of units across waveform durations for each saccadic modulation response type (early, late, suppressive). Grey in the histograms represents the full population with color representing significant saccade tuning to blank images C) The distribution of early, late, and suppressive response types as a function of spike duration and cortical recording depth (types represented by color). D) Scatter plot showing the distribution of narrow and broad spiking units (yellow and green) across cortical depth as a function of saccade modulation peak latency. Filled circles represent units with significant saccade tuning on blank images. E) Mean normalized saccadic modulation separated out by narrow spiking units (yellow) and broad spiking units (green) across Superficial (Top), Input (Middle), and Deep (Bottom) layers. Solid lines show the modulation on natural images, while dashed lines show the modulation on blank screens, error bars are 2 sem. F,G) Histograms showing spike waveform durations for units flagged for having saccadic omission (black) and early response units with significant saccade direction tuning in the dark (red). H) The distribution of spike waveform duration as a function of cortical depth for saccadic omission and early response units with saccade tuning in the dark.

The distribution of spike durations and laminar position also varied based on the saccade response type, with early and late response types showing a higher proportion of narrow spikes and occurring more in the input and deep layers whereas suppressive response types showed broader spikes and appeared more often in superficial layers (**Figure 7B**). The early and late populations differed significantly in spike duration from the suppressive population (early vs. suppressive, kstest, p < 0.001; late vs. suppressive, kstest, p < 0.001) because they had more narrow than broad spiking units while suppressive populations had predominantly broad spiking waveforms (Early: 46% Narrow, 41% Broad; Late: 45% Narrow, 38%; Suppressive: Broad 28% Narrow, 58% Broad). These differences were also evident plotting each of the response type’s spike duration and cortical depth (**Figure 7C**). Early and late response types (orange and magenta) were sampled more frequently in the input layer where there was a higher proportion of narrow spikes (Input Layer Proportion: Early 39.7%, Late 46.3%, Suppressive 30.3%) which differed significantly from suppressive units (Proportion Test: Early vs. Late, p=0.071; Early vs. Supp., p= 0.006; Late vs Supp, p<0.001). Suppressive units also differed by appearing more in the superficial layers than early or late units (Superficial Layer Proportion: Early 16.7%, Late 8.5%, Suppressive 26.2%) and differed in their proportion significantly (Proportion Test: Early vs. Late, p = 0.001; Early vs. Supp., p= 0.001; Late vs Supp, p<0.001). The proportion of response types was relatively uniform in the deep layers (Deep Layer Proportion: Early 44.0%, Late 45.1%, Suppressive 43.8%). In brief, these laminar distinctions suggest a disassociation between early response types in input/deep layers and suppressive responses in superficial.

We also found differences in the dynamics of inhibitory and excitatory cell activity across layers with inhibition leading excitation (**Figure 7D**). To better visualize the dynamics of the inhibitory and excitatory populations, we plotted the mean modulation per layer split by narrow and broad spiking categories (**Figure 7E**). In all layers the narrow spiking neurons showed the strongest and earliest saccadic modulation. These differences reached significance in the 50 ms following saccade onset in the deep layer but were not yet significant in the superficial and input layers (Superficial: Narrow: 1.276, Broad: 1.206, p=0.814; Input: Narrow: +1.637, Broad: +1.394, rank-sum, p=0.583; Deep: Narrow: +1.499, Broad: +1.291, rank-sum, p=0.024). But after 50ms there was stronger modulation among narrow spiking units for the input and superficial layers (Superficial: Narrow: +1.379, Broad: +1.002, rank-sum, p < 0.001, **Figure7C:Top;** Input layer: Narrow: +2.160, Broad: +1.481, rank-sum, p < 0.001, **Figure 7C:Middle**), while the the deep layer was no longer significantly different (Narrow: +1.640, Broad: +1.670, rank-sum, p = 0.764) (**Figure 7C:Bottom**). Overall, narrow spiking units led the response to saccades, predominantly with a positive increase in rate over baseline rather than with suppression, and as such, could have driven suppression in the rest of the population.

Last, we found that two interesting subsets of units discussed previously, early response units that respond in the dark (Early Dark Units) and units that omit a motion response when caused by real saccades (Saccadic Omission Units) were differentially distributed across layers and cell types. For the “Early Dark” units, we found the population had a bias towards narrow spike waveforms (52.8% Narrow, 39.6% Broad) (**Figure 7D**). By contrast, the saccadic omission neurons showed the opposite bias towards broad spike waveforms (20% Narrow, 73.8% Broad) (**Figure 7E**), which was significantly different (test of proportions, p=0.0002). Overlaying these two subsets as a scatter plot across spike duration and cortical depth, we found that Early Dark units were on average in the input and deep layers while the Omission Units were on average biased towards superficial layers (**Figure 7F**). The proportion of neurons in the superficial layers for Omission Units was 51.2% as compared to only 20.8% among Early Dark units (test of proportions, p = 0.0008). These results support that there are distinctions within MT/MTC neural populations wherein early response units support fast latency detection of saccades while other neurons responding at slower latencies are able to omit motion induced responses for real saccades, and that these types differ by laminar position and cell type.

## Discussion

We investigated how saccade eye movements modulate neural activity in areas MT/MTC using free-viewing behavioral paradigms with saccades made on natural images, blank images, and dark screens. We found a variety of responses that formed a continuum from excitatory early responses to slower responses and purely suppressive responses. We sought to disassociate if visual or extra-retinal cues drove the observed saccadic modulations. We found that over a third of the population had early positive responses that were tuned for the direction of the saccade, and that this tuning persisted for saccades on blank screens or in complete darkness suggesting some role for an extra-retinal (corollary discharge) input. But these neurons also responded robustly when visual motion that simulated saccades were shown in the absence of an eye movement, and in that condition retained consistent direction tuning. Other neurons that exhibited late or purely suppressive responses were not well tuned for saccade direction. More so, some neurons responded strongly to simulated saccades but had negligible responses to the motion induced by real saccades. That subset of neurons was consistent with a previous report from macaque area MT and MST (Thiele et al., 2002) that found neurons which suppress motion responses that are related to self-induced motion. Those ‘saccadic omission” neurons may underlie the neural correlate of perceptual stability during saccades.

One motivation of the present work was to compare the dynamics of saccade modulation against an earlier study in V1 that found dynamics consistent with a coarse-to-fine processing strategy (Parker et al., 2023). We found that units with early or late positive responses were tuned for visual stimuli with lower spatial frequencies while those with suppressive responses preferred higher spatial frequencies (**Figure 2C,D**). This could support a coarse-to-fine processing strategy similar to V1. However, there were some distinctions in our distribution from the V1 study. The proportion of neurons in MT/MTC with early response properties was much higher than V1, reaching 39% of the population as compared to 19% in V1 (Parker et al., 2023). Further, the dominant response in V1 was bi-phasic with suppression followed by an excitatory rebound, which was relatively rare in MT/MTC. These differences may reflect a distinction between magnocellular (fast, low resolution) and parvocellular (slow, high resolution) input pathways to V1, whereas areas MT/MTC receive predominantly magnocellular input (Maunsell et al., 1990). Taken together, early responses appear to be more characteristic of magnocellular pathway input, which would explain why they are stronger in MT/MTC than V1.

Our findings on the laminar and spike width distribution were broadly consistent with a previous laminar study of saccadic modulation in macaque area V4 (Denagamage et al., 2023). Laminar and cell type distinctions can provide insights into the origin of saccade modulation. For example, information fed back from frontal oculomotor areas terminates in superficial and deep layers of visual cortex (Anderson et al., 2011; Larkum, 2013; Markov et al., 2013) whereas feedback from the superior colliculus through the pulvinar predominately goes to input layers (Berman & Wurtz, 2010; Trojanowski & Jacobson, 1977).The previous study in V4 reported that narrow spiking (putative inhibitory) neurons in the input layer had an excitatory early response to saccades while broad spiking (putative excitatory) units were generally suppressed. It was hypothesized that inhibitory neurons in the input layer might receive corollary discharge signals, potentially relayed back from the SC through the pulvinar which projects to input layers. Those early responding inhibitory cells could then suppress saccade induced activity among excitatory cells in other layers. Our findings are somewhat consistent with this hypothesis in that the early response units in our population are biased towards being narrow spiking neurons and do show higher sampling in input layers although also in deep layers (**Figure 7D,E)**. Likewise, those neurons in our dataset exhibiting suppressive responses were biased towards being broad spiking and occurred more frequently in superficial layers (**Figure 7B,C**). Thus, our findings would generally support a similar circuit model in which the detection of saccades occurs among inhibitory units in input and deep layers and drives suppression in broad spiking units in superficial layers. It could be that excitatory neurons in superficial layers, the putative projection neurons relaying information to downstream areas, would then better support the perception of stability during saccades.

While our study highlights distinctions between layers and cell types, there are also some limitations. First, the accuracy with which we can localize the input layer from the CSD analysis has some uncertainty in its measurement, at least on the order of 50-100 microns. Further, the CSD is thought to detect the bottom of the input layer both in V1 and MT (Schroeder et al., 1998), and we have fixed the top of input layer to be 300 microns above the bottom. This assumes a fixed width of the input layer based on histology from marmoset MT (Bourne et al., 2007). But our laminar recordings also compress cortical tissue during penetration through the layers, which is likely to vary between recordings, and would add variability to our measurements. Thus, there are uncontrolled sources of variation in our laminar estimates that would be expected to obscure any real laminar distinctions.

Despite those limitations we did find differences between the responses across layers in MT/MTC. Specifically, we found that there were a significantly higher proportion of narrow spiking units localized to the input layer. Nearly half of the neurons in input layer had narrow spikes (**Figure 7A**), a value that would not fit the known distribution of fast spiking interneurons in MT or other parts of visual cortex (Disney et al., 2014; Hendry et al., 1987) In area V1 it is known that there are excitatory neurons in the input layer that express Kv3.1b potassium channels resulting in narrow spike waveforms (Kelly & Hawken, 2020). It is unknown if similar excitatory cells like those might also be present in the input layer of MT, but there is clear evidence that area MT shows distinctions in the cellular composition of its input layer (Bourne et al., 2007). Thus, while our laminar and cell type distributions are roughly consistent with the former V4 study (Denagamage et al., 2023), some observed differences are likely due to real differences.

Several previous studies have argued that oculomotor feedback is involved in driving saccadic suppression in area MT (Berman & Wurtz, 2010; Trojanowski & Jacobson, 1977), but the origins of suppression remain far from definitive. Anatomical pathways stemming from motor planning regions such as the Frontal Eye Fields or the Superior Colliculus could provide corollary signals to visual areas like MT to trigger suppression (Brenner et al., 2023; Duffy & Lombroso, 1968; Holt, 1903; Matin, 1974; Miura & Scanziani, 2022; Zuber et al., 1966). It is also possible that suppression could be imposed in early processing stages and inherited by area MT. For example, stimulation of the saccade generator has been shown to suppress activity in the lateral geniculate nucleus (LGN) demonstrating a causal extra-retinal effect (Doty et al., 1973) and saccades in the dark suppress LGN activity (Bartlett et al., 1976; Lee & Malpeli, 1998; Royal et al., 2006) as well as V1 activity (Duffy & Burchfiel, 1975; Kagan et al., 2008; Kayama et al., 1979; McFarland et al., 2015). Thus, extra-retinal inputs could impact suppression at earlier stages of processing to influence the subsequent processing in MT/MTC.

However, it has also been argued that the retinal motion itself, generated by the saccade, could trigger suppression as a masking effects (Beeler, 1967; Campbell & Wurtz, 1978; Mackay, 1970; Sperling, 1990). Recent work suggests that saccadic suppression can even start as early as the retina where there are no feedback signals (Idrees et al., 2020), and propose instead that corollary discharge might operate at later stages of processing by speeding recovery from suppression (Diamond et al., 2000; Idrees et al., 2020). There are also visual neurons that are known to detect saccades based on wide-field motion signals. Neurons termed “jerk” cells are found in the pretectum of both cats and primates that are selective to fast wide-field motion (velocities above 100 degrees/sec), that have large receptive fields, and that are highly sensitive to saccades (Schweigart and Hoffmann, 1992; Schmidt, 1996; Sudkamp and Schmidt, 2000; Price and Ibbotson, 2001). These neurons also respond to saccades in the dark, though with much slower latencies than their visually driven responses (Schweigart and Hoffman, 1992; Schmidt, 1996). These regions of pretectum containing “jerk” cells project to extra-striate visual cortex by pathways running through the pulvinar (Ugolini & Graf, 2024), and thus could provide inputs to MT/MTC within pathways overlapping feedback from the superior colliculus.

Our studies support that both extra-retinal and visual cues are integrated to drive early saccade tuned responses in MT/MTC, with visual signals providing the dominant of these signals. Early cells responded both to real saccades on natural images, as well as saccades on blank screens or to saccades in complete darkness, and did so with consistent tuning in saccade direction across those conditions. In support of the visual origin of their response, they responded with similar tuning for simulated saccades (**Figure 6B,E**). Supporting some role for an extra-retinal signal among these cells, we found that many of them retained a significant response in the dark and that it had consistent directional tuning with saccades in the light (**Figure 5C,D**).

However, in the dark these neurons had much slower response latencies that peaked after 50 ms from saccade onset as opposed to earlier than 50 ms in light. This aspect of their responses resembles that of “jerk” neurons reported from the pretectum, which respond at slower latencies to saccades in the dark (Schmidt, 1996). However, this slow latency response could also reflect residual low light responses in the dark due to scotopic vision, which would be mediated through the rod visual pathways (Conner & MacLeod, 1977; McKyton et al., 2024; Vaughan et al., 1966). We consider this possibility unlikely though because the saccade response in darkness did not strengthen over time with increased dark adaptation (**Supplementary Figure 3)**. Thus, we do find support for an extra-retinal contribution to MT/MTC responses, although it does not appear to be a strongest input and normally would complement a visual signal that supports faster saccade detection.

A final key finding in the present study was that a subset of neurons in the population omit their responses to motion induced by real but not simulated saccades. This subset is consistent with an earlier report from MT and MST using motion stimuli that simulated saccades and found that roughly 20% of the population discounted saccade-induced retinal motion (Thiele et al., 2002). Beyond the earlier study, we found that these ‘saccadic omission’ neurons sampled preferentially from broad spiking (putative excitatory) neurons and that they were biased towards superficial layers of cortex (**Figure 6F,G**). These neurons would thus overlap with the projection neurons from areas MT/MTC that relay motion information to higher cortical areas, and thus could better represent the perceptual effects of saccadic omission. These findings also support that there is a circuit in MT/MTC both for detecting and discounting saccadic motion, but additional work will be necessary to determine the mechanisms at a cellular level. Recent advances for targeting specific cell classes in non-human primates(De et al., 2020; El-Shamayleh & Horwitz, 2019; Federer et al., 2024) as well as specific projection pathways (Shaw et al., 2026) holds promise to make these kinds of circuit manipulations possible.

## Acknowledgments

would like to thank Dina Graf and members of the Mitchell lab for help with marmoset care and handling. This work was supported by NIH grants R01 EY030998 (JFM, SC, AB), NIH T32EY007125 (AB) and T32 EY007125 (SC).

## Bibliography

Anderson, J. C., Kennedy, H., & Martin, K. A. C. (2011). Pathways of Attention: Synaptic Relationships of Frontal Eye Field to V4, Lateral Intraparietal Cortex, and Area 46 in Macaque Monkey. Journal of Neuroscience, 31(30), 10872–10881. 10.1523/JNEUROSCI.0622-11.2011

Baloh, R. W., Sills, A. W., Kumley, W. E., & Honrubia, V. (1975). Quantitative measurement of saccade amplitude, duration, and velocity. Neurology, 25(11), 1065–1065. 10.1212/WNL.25.11.1065

Bartlett, J. R., Doty, R. W., Lee, B. B., & Sakakura, H. (1976). Influence of saccadic eye movements on geniculostriate excitability in normal monkeys. Experimental Brain Research, 25(5), 487–509. 10.1007/BF00239783

Beeler, G. W. (1967). Visual threshold changes resulting from spontaneous saccadic eye movements. Vision Research, 7(9), 769–775. 10.1016/0042-6989(67)90039-9

Berman, R. A., Cavanaugh, J., McAlonan, K., & Wurtz, R. H. (2017). A circuit for saccadic suppression in the primate brain. Journal of Neurophysiology, 117(4), 1720–1735. 10.1152/jn.00679.2016

Berman, R. A., & Wurtz, R. H. (2010). Functional Identification of a Pulvinar Path from Superior Colliculus to Cortical Area MT. Journal of Neuroscience, 30(18), 6342–6354. 10.1523/JNEUROSCI.6176-09.2010

Berman, R. A., & Wurtz, R. H. (2011). Signals Conveyed in the Pulvinar Pathway from Superior Colliculus to Cortical Area MT. Journal of Neuroscience, 31(2), 373–384. 10.1523/JNEUROSCI.4738-10.2011

Bourne, J. A., Warner, C. E., Upton, D. J., & Rosa, M. G. P. (2007). Chemoarchitecture of the middle temporal visual area in the marmoset monkey (Callithrix jacchus): Laminar distribution of calcium-binding proteins (calbindin, parvalbumin) and nonphosphorylated neurofilament. Journal of Comparative Neurology, 500(5), 832–849. 10.1002/cne.21190

Brainard, D. H. (1997). The Psychophysics Toolbox. Spatial Vision, 10(4), 433–436. 10.1163/156856897X00357

Bremmer, F., Kubischik, M., Hoffmann, K.-P., & Krekelberg, B. (2009). Neural Dynamics of Saccadic Suppression. The Journal of Neuroscience, 29(40), 12374–12383. 10.1523/JNEUROSCI.2908-09.2009

Brenner, J. M., Beltramo, R., Gerfen, C. R., Ruediger, S., & Scanziani, M. (2023). A genetically defined tecto-thalamic pathway drives a system of superior-colliculus-dependent visual cortices. Neuron, 111(14), 2247–2257.e7. 10.1016/j.neuron.2023.04.022

Bucklaew, A., Coop, S. H., & Mitchell, J. F. (2023). Electrophysiology of Laminar Cortical Activity in the Common Marmoset. Journal of Visualized Experiments, (198). 10.3791/65397

Campbell, F. W., & Wurtz, R. H. (1978). Saccadic omission: Why we do not see a grey-out during a saccadic eye movement. Vision Research, 18(10), 1297–1303. 10.1016/0042-6989(78)90219-5

Castet, E., Jeanjean, S., & Masson, G. S. (2002). Motion perception of saccade-induced retinal translation. Proceedings of the National Academy of Sciences, 99(23), 15159–15163. 10.1073/pnas.232377199

Castet, E., & Masson, G. S. (2000). Motion perception during saccadic eye movements. Nature Neuroscience, 3(2), Article 2. 10.1038/72124

Chen, C.-Y., & Hafed, Z. M. (2017). A neural locus for spatial-frequency specific saccadic suppression in visual-motor neurons of the primate superior colliculus. Journal of Neurophysiology, 117(4), 1657–1673. 10.1152/jn.00911.2016

Chukoskie, L., & Movshon, J. A. (2009). Modulation of Visual Signals in Macaque MT and MST Neurons During Pursuit Eye Movement. Journal of Neurophysiology, 102(6), 3225–3233. 10.1152/jn.90692.2008

Conner, J. D., & MacLeod, D. I. (1977). Rod photoreceptors detect rapid flicker. Science, 195(4279), 698–699. 10.1126/science.841308

Connors, B. W., & Gutnick, M. J. (1990). Intrinsic firing patterns of diverse neocortical neurons. Trends in Neurosciences, 13(3), 99–104. 10.1016/0166-2236(90)90185-D

Coop, S. H., Crutcher, G. W., Abrham, Y. T., Bucklaew, A., & Mitchell, J. F. (2024). Pre-saccadic enhancement of target stimulus motion influences post-saccadic smooth eye movements (p. 2022.10.10.511640). bioRxiv. 10.1101/2022.10.10.511640

De, A., El-Shamayleh, Y., & Horwitz, G. D. (2020). Fast and reversible neural inactivation in macaque cortex by optogenetic stimulation of GABAergic neurons. eLife, 9, e52658. 10.7554/eLife.52658

Denagamage, S., Morton, M. P., Hudson, N. V., Reynolds, J. H., Jadi, M. P., & Nandy, A. S. (2023). Laminar mechanisms of saccadic suppression in primate visual cortex. Cell Reports, 42(7). 10.1016/j.celrep.2023.112720

Diamond, M. R., Ross, J., & Morrone, M. C. (2000). Extraretinal Control of Saccadic Suppression. The Journal of Neuroscience, 20(9), 3449–3455. 10.1523/JNEUROSCI.20-09-03449.2000

Disney, A. A., Alasady, H. A., & Reynolds, J. H. (2014). Muscarinic acetylcholine receptors are expressed by most parvalbumin-immunoreactive neurons in area MT of the macaque. Brain and Behavior, 4(3), 431–445. 10.1002/brb3.225

Doty, R. W., Wilson, P. D., Bartlett, J. R., & Pecci-Saavedra, J. (1973). Mesencephalic control of lateral geniculate nucleus in primates. I. Electrophysiology. Experimental Brain Research, 18(2), 189–203. 10.1007/BF00234723

Duffy, F. H., & Burchfiel, J. (1975). Eye movement-related inhibition of primate visual neurons. Brain Research, 89(1), 121–132. 10.1016/0006-8993(75)90139-0

Duffy, F. H., & Lombroso, C. T. (1968). Electrophysiological Evidence for Visual Suppression prior to the Onset of a Voluntary Saccadic Eye Movement. Nature, 218(5146), 1074–1075. 10.1038/2181074a0

El-Shamayleh, Y., & Horwitz, G. D. (2019). Primate optogenetics: Progress and prognosis. Proceedings of the National Academy of Sciences, 116(52), 26195–26203. 10.1073/pnas.1902284116

Engbert, R., & Mergenthaler, K. (2006). Microsaccades are triggered by low retinal image slip. Proceedings of the National Academy of Sciences, 103(18), 7192–7197. 10.1073/pnas.0509557103

Federer, F., Balsor, J., Ingold, A., Babcock, D. P., Dimidschstein, J., & Angelucci, A. (2024). Laminar specificity and coverage of viral-mediated gene expression restricted to GABAergic interneurons and their parvalbumin subclass in marmoset primary visual cortex. eLife, 13, RP97673. 10.7554/eLife.97673

Fischer, B., & Weber, H. (1993). Express saccades and visual attention. Behavioral and Brain Sciences, 16(3), 553–567. 10.1017/S0140525X00031575

Hendry, S. H. C., Schwark, H. D., Jones, E. G., & Yan, J. (1987). Numbers and Proportions of GABA-lmmunoreactive Neurons in Different Areas of Monkey Cerebral Cortex. The Journal of Neuroscience, 7(5), 1503–1519.

Holt, E. B. (1903). Eye-movement and central anaesthesia. The Psychological Review: Monograph Supplements, 4(1), 1–45.

Idrees, S., Baumann, M. P., Franke, F., Münch, T. A., & Hafed, Z. M. (2020). Perceptual saccadic suppression starts in the retina. Nature Communications, 11(1), 1977. 10.1038/s41467-020-15890-w

Kagan, I., Gur, M., & Snodderly, D. M. (2008). Saccades and drifts differentially modulate neuronal activity in V1: Effects of retinal image motion, position, and extraretinal influences. Journal of Vision, 8(14), 19. 10.1167/8.14.19

Kayama, Y., Riso, R. R., Bartlett, J. R., & Doty, R. W. (1979). Luxotonic responses of units in macaque striate cortex. Journal of Neurophysiology, 42(6), 1495–1517. 10.1152/jn.1979.42.6.1495

Kelly, J. G., & Hawken, M. J. (2020). GABAergic and non-GABAergic subpopulations of Kv3.1b-expressing neurons in macaque V2 and MT: Laminar distributions and proportion of total neuronal population. Brain Structure and Function, 225(3), 1135–1152. 10.1007/s00429-020-02065-y

Kwon, S., Rolfs, M., & Mitchell, J. F. (2019). Presaccadic motion integration drives a predictive postsaccadic following response. Journal of Vision, 19(11), 12. 10.1167/19.11.12

Larkum, M. (2013). A cellular mechanism for cortical associations: An organizing principle for the cerebral cortex. Trends in Neurosciences, 36(3), 141–151. 10.1016/j.tins.2012.11.006

Lee, D., & Malpeli, J. G. (1998). Effects of Saccades on the Activity of Neurons in the Cat Lateral Geniculate Nucleus. Journal of Neurophysiology, 79(2), 922–936. 10.1152/jn.1998.79.2.922

Leopold, D. A., & Logothetis, N. K. (1998). Microsaccades differentially modulate neural activity in the striate and extrastriate visual cortex. Experimental Brain Research, 123(3), 341–345. 10.1007/s002210050577

Mackay, D. M. (1970). Elevation of Visual Threshold by Displacement of Retinal Image. Nature, 225(5227), 90–92. 10.1038/225090a0

Markov, N. T., Ercsey-Ravasz, M., Van Essen, D. C., Knoblauch, K., Toroczkai, Z., & Kennedy, H. (2013). Cortical High-Density Counterstream Architectures. Science, 342(6158), 1238406. 10.1126/science.1238406

Matin, E. (1974). Saccadic suppression: A review and an analysis. Psychological Bulletin, 81(12), 899–917. 10.1037/h0037368

Maunsell, J., Nealey, T., & DePriest, D. (1990). Magnocellular and parvocellular contributions to responses in the middle temporal visual area (MT) of the macaque monkey. The Journal of Neuroscience, 10(10), 3323–3334. 10.1523/JNEUROSCI.10-10-03323.1990

McFarland, J. M., Bondy, A. G., Saunders, R. C., Cumming, B. G., & Butts, D. A. (2015). Saccadic modulation of stimulus processing in primary visual cortex. Nature Communications, 6(1), 8110. 10.1038/ncomms9110

McKyton, A., Elul, D., & Levin, N. (2024). Seeing in the dark: High-order visual functions under scotopic conditions. iScience, 27(2), 108929. 10.1016/j.isci.2024.108929

Mitchell, J. F., Reynolds, J. H., & Miller, C. T. (2014). Active Vision in Marmosets: A Model System for Visual Neuroscience. Journal of Neuroscience, 34(4), 1183–1194. 10.1523/JNEUROSCI.3899-13.2014

Mitchell, J. F., Sundberg, K. A., & Reynolds, J. H. (2007). Differential Attention-Dependent Response Modulation across Cell Classes in Macaque Visual Area V4. Neuron, 55(1), 131–141. 10.1016/j.neuron.2007.06.018

Mitzdorf, U. (1985). Current source-density method and application in cat cerebral cortex: Investigation of evoked potentials and EEG phenomena. Physiological Reviews, 65(1), 37–100. 10.1152/physrev.1985.65.1.37

Miura, S. K., & Scanziani, M. (2022). Distinguishing externally from saccade-induced motion in visual cortex. Nature, 610(7930), 135–142. 10.1038/s41586-022-05196-w

Nummela, S. U., Coop, S. H., Cloherty, S. L., Boisvert, C. J., Leblanc, M., & Mitchell, J. F. (2017). Psychophysical measurement of marmoset acuity and myopia. Developmental Neurobiology, 77(3), 300–313. 10.1002/dneu.22467

Parker, P. R. L., Martins, D. M., Leonard, E. S. P., Casey, N. M., Sharp, S. L., Abe, E. T. T., Smear, M. C., Yates, J. L., Mitchell, J. F., & Niell, C. M. (2023). A dynamic sequence of visual processing initiated by gaze shifts. Nature Neuroscience, 26(12), 2192–2202. 10.1038/s41593-023-01481-7

Paxinos, G. (2012). The marmoset brain in stereotaxic coordinates (1st ed). Academic Press.

Pelli, D. G. (1997). The VideoToolbox software for visual psychophysics: Transforming numbers into movies. Spatial Vision, 10(4), 437–442. 10.1163/156856897x00366

Pierrot-Deseilligny, C., Rivaud, S., Gaymard, B., Müri, R., & Vermersch, A.-I. (1995). Cortical control of saccades. Annals of Neurology, 37(5), 557–567. 10.1002/ana.410370504

Rosa, M. G. P., & Elston, G. N. (1998). Visuotopic organisation and neuronal response selectivity for direction of motion in visual areas of the caudal temporal lobe of the marmoset monkey (Callithrix jacchus): Middle temporal area, middle temporal crescent, and surrounding cortex. Journal of Comparative Neurology, 393(4), 505–527. 10.1002/(SICI)1096-9861(19980420)393:4%3C505::AID-CNE9%3E3.0.CO;2-4

Royal, D. W., Sáry, Gy., Schall, J. D., & Casagrande, V. A. (2006). Correlates of motor planning and postsaccadic fixation in the macaque monkey lateral geniculate nucleus. Experimental Brain Research, 168(1), 62–75. 10.1007/s00221-005-0093-z

Schmidt, M. (1996). Neurons in the cat pretectum that project to the dorsal lateral geniculate nucleus are activated during saccades. Journal of Neurophysiology, 76(5), 2907–2918. 10.1152/jn.1996.76.5.2907

Schroeder, C. E., Mehta, A. D., & Givre, S. J. (1998). A spatiotemporal profile of visual system activation revealed by current source density analysis in the awake macaque. Cerebral Cortex, 8(7), 575–592. 10.1093/cercor/8.7.575

Shaw, L., Padmanabhan, K., Bucklaew, A., Mitchell, J. F., & Wang, K. H. (2026). Projection-specific intersectional optogenetics for precise excitation and inhibition in the marmoset brain. Cell Reports Methods, 6(4). 10.1016/j.crmeth.2026.101368

Sperling, G. (1990). Comparison of perception in the moving and stationary eye. Reviews of Oculomotor Research, 4, 307–351.

Thiele, A., Henning, P., Kubischik, M., & Hoffmann, K.-P. (2002). Neural Mechanisms of Saccadic Suppression. Science, 295(5564), 2460–2462. 10.1126/science.1068788

Trojanowski, J. Q., & Jacobson, S. (1977). The morphology and laminar distribution of cortico-pulvinar neurons in the Rhesus monkey. Experimental Brain Research, 28(1), 51–62. 10.1007/BF00237085

Ugolini, G., & Graf, W. (2024). Pathways from the superior colliculus and the nucleus of the optic tract to the posterior parietal cortex in macaque monkeys: Functional frameworks for representation updating and online movement guidance. European Journal of Neuroscience, 59(10), 2792–2825. 10.1111/ejn.16314

Vaughan, H. G., Costa, L. D., & Gilden, L. (1966). The functional relation of visual evoked response and reaction time to stimulus intensity. Vision Research, 6(11), 645–656. 10.1016/0042-6989(66)90076-9

Volkmann, F. C. (1986). Human visual suppression. Vision Research, Twenty-Fifth Anniversary Issue Of, 26(9), 1401–1416. 10.1016/0042-6989(86)90164-1

Wolfe, J. M., Alvarez, G. A., Rosenholtz, R., Kuzmova, Y. I., & Sherman, A. M. (2011). Visual search for arbitrary objects in real scenes. Attention, Perception & Psychophysics, 73(6), 1650– 1671. 10.3758/s13414-011-0153-3

Yates, J. L., Coop, S. H., Sarch, G. H., Wu, R.-J., Butts, D. A., Rucci, M., & Mitchell, J. F. (2023). Detailed characterization of neural selectivity in free viewing primates. Nature Communications, 14(1), 3656. 10.1038/s41467-023-38564-9

Zuber, B. L., Stark, L., & Lorber, M. (1966). Saccadic suppression of the pupillary light reflex. Experimental Neurology, 14(3), 351–370. 10.1016/0014-4886(66)90120-8

