## Supplementary Text and Figures for "Neural subpopulations in marmoset area MT/MTC detect and discount saccade-related retinal motion"

Neural subpopulations in marmoset area MT/MTC show diverse saccade-related modulation

**Results:**

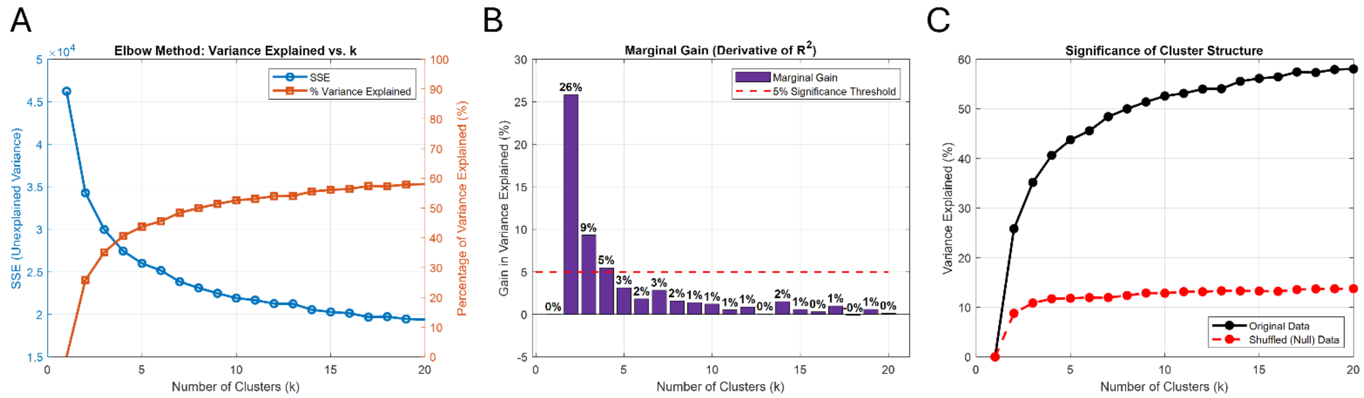


**Supplemental Figure 1: Finding Optimal K Value** (A) The total within-cluster sum of squares (SSE) dropped with increasing numbers of fit clusters (k), shown in blue, while the percentage variance explained by the SSE increased over the same range (orange), showing a cross-over just after k=3. (B) The Marginal Gain of each k value plotted against the Heuristic Threshold of 5% (red dashed line) supported an optimal k of 3. (C) The original data (black) showed larger variance explained compared to a shuffled version of the data (red).

To objectively determine the functional categories of saccadic modulation across the MT/MTC population, we performed a k-means clustering on the neural response profiles. Plotting the total within-cluster sum of squared errors (SSE) alongside the percentage of variance explained across a range of cluster values revealed a cross of the SSE and Variance Explained after k = 3 (**Supplementary Figure 1A**). This choice was further confirmed by calculating the marginal gain in variance explained for each additional cluster, which demonstrated that k =3 was the optimal threshold before falling below our 5% heuristic cutoff (**Supplementary Figure 1B**). Finally, comparing the variance explained in our original dataset against a shuffled control confirmed that these three functional clusters represent true biological structure rather than random variation in the neural recordings (**Supplementary Figure 1C**).

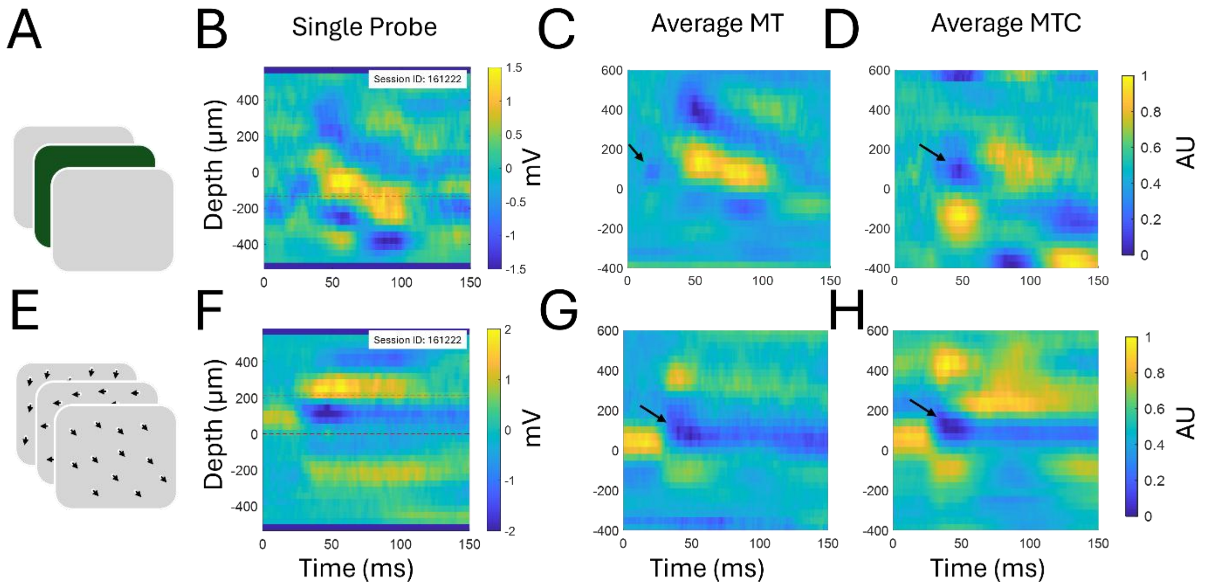


**Supplemental Figure 2: Example Current Source Density Analysis** (A) Full Field Flashing Stimuli were used to probe the location of input layer across recording shanks (B) Single Session example CSD (C,D,G,H) Average CSDs across sessions were generated by first normalizing each session CSD between 0 and 1, where below 0.5 reflects a sink and above a source, and then aligning them vertically to the identified sink before averaging. (C) Average MT CSD shown with the black arrow denoting the location of the current sink (D) Average MTC CSD (E) Full field Motion Stimuli that translated with 100% coherence for 400 ms with an interstimulus interval of 400 ms were used to evoke a CSD response. CSD analyses were time-locked to motion onset. (F) Single Shank Example CSD (G) Average MT CSD shown with black arrow denoting the location of the current sink (H) Average MTC CSD for the motion onset.

To map the laminar profile of our recordings and identify the input layer (Layer 4) across area MT and MTC, we performed Current Source Density (CSD) analysis using two distinct visual stimuli. First, full-field flashing stimuli were presented to elicit sharp, early sink-source transitions, yielding clear single-session profiles as well as population-averaged CSDs for both MT and MTC where the primary early current sink was marked (**Supplementary Figure 2A-D**). Second, 100% coherent full-field motion stimuli were used as an independent visual probe (see methods for stimulus details and timing), similarly revealing consistent single-shank CSD profiles and population averages (**Supplementary Figure 2E-H**). The early current sinks identified under both conditions aligned closely, providing a robust baseline for localized laminar assignments across both cortical areas.

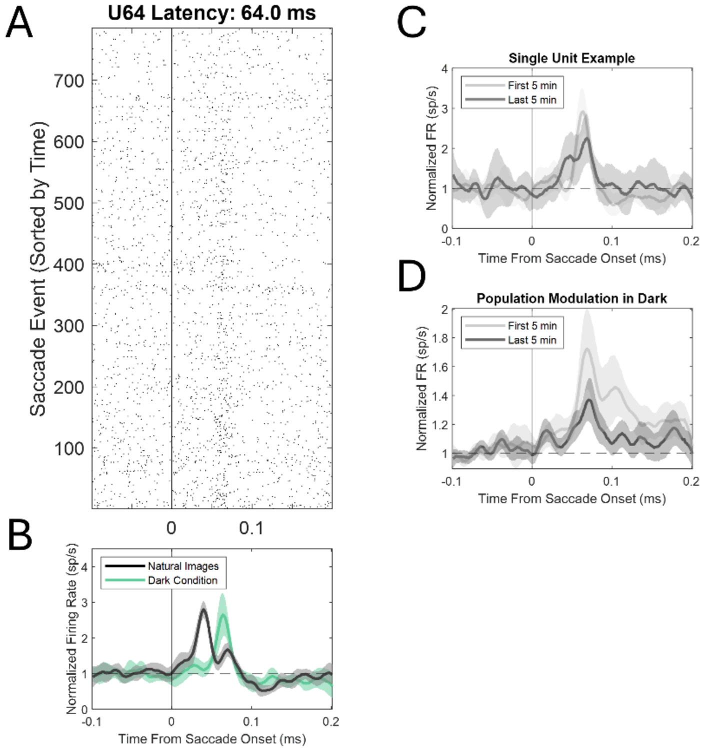


**Supplemental Figure 3: Dark Condition Modulation Over Time** (A-B) Single unit example showing a PSTH (Top) and mean normalized response (bottom) of an early unit showing modulation in natural image condition (black) and in the dark condition (green). (C) A single unit example of saccade-related modulation in the dark for the first 5 minutes (light grey) and last 5 minutes (black) of dark adaptation. (D) Average modulation from 20 example units having strong saccadic modulation in the dark, separating the modulation during the first 5 minutes of the dark condition (light grey) and the last 5 minutes of the dark condition (dark grey).

To ensure that saccade-related modulation recorded in complete darkness reflected stable electrical signals rather than transient dark adaptation or conversely arousal artifacts, we tracked single-unit and population responses across time. An example of an individually significant early-responding unit in the dark showed a significant difference in its response peak for saccades in the light viewing natural images (black) as compared to in darkness (green), though with little different in percentage modulate relative to baseline (**Supplementary Figure 3A-B**). To consider the possibility that the slower latency dark responses could reflect a response driven by scoptic visual inputs, we assessed if the response varied with the duration of dark adaptation by comparing the average responses in the first 5 minutes of being in the darkness versus the last 5 minutes (these blocks last on average 15 minutes). A single unit example showed little difference between these different time intervals, suggesting dark adaptation did not play a key role (**Supplementary Figure 3C**). Across the population, comparing the average modulation during the first 5 minutes of dark exposure to the final 5 minutes revealed similar response profiles, with a trend for a stronger response (light gray) in the first 5 minutes as compared to the last five minutes (black), supporting rather that arousal may have diminished with prolonged time in the dark as opposed to adaptation under scotopic vision (**Supplementary Figure 3D**).
